# Distinct Aurora B pools at the inner centromere and kinetochore have different contributions to meiotic and mitotic chromosome segregation

**DOI:** 10.1101/2023.02.05.527197

**Authors:** Gisela Cairo, Cora Greiwe, Gyu Ik Jung, Cecilia Blengini, Karen Schindler, Soni Lacefield

## Abstract

Proper chromosome segregation depends on establishment of bioriented kinetochore-microtubule attachments, which often requires multiple rounds of release and reattachment. Aurora B and C kinases phosphorylate kinetochore proteins to release tensionless attachments. Multiple pathways recruit Aurora B/C to the centromere and kinetochore. We studied how these pathways contribute to anaphase onset timing and correction of kinetochore-microtubule attachments in budding yeast meiosis and mitosis. We find that the pool localized by the Bub1/Bub3 pathway sets the normal duration of meiosis and mitosis, in differing ways. Our meiosis data suggests that disruption of this pathway leads to PP1 kinetochore localization, which dephosphorylates Cdc20 for premature anaphase onset. For error correction, the Bub1/Bub3 and COMA pathways are individually important in meiosis but compensatory in mitosis. Finally, we find that the haspin and Bub1/3 pathways function together to ensure error correction in mouse oogenesis. Our results suggest that each recruitment pathway localizes spatially distinct kinetochore-localized Aurora B/C pools that function differently between meiosis and mitosis.

## Introduction

Faithful partitioning of genetic material during mitosis and meiosis is fundamental to survival of an organism. Errors in chromosome segregation can result in aneuploidy, or a gain or loss of chromosomes. Aneuploidy during mitosis is prevalent in tumor cells and is likely a major factor driving tumor evolution (Funk et al., 2016). In meiosis, aneuploidy can lead to trisomy conditions, miscarriage, and infertility (Nagaoka et al., 2012). To ensure that chromosomes segregate faithfully, spindle microtubules must properly attach to kinetochores, a large protein complex assembled on centromeric DNA (Cairo and Lacefield, 2020). During mitosis, DNA replication is followed by the attachment of sister chromatid kinetochores to microtubules emanating from opposite spindle poles, a state known as biorientation. The bipolar attachments result in kinetochores that are under tension due to the poleward forces of the spindle resisted by cohesins that encircle the sister chromatids. At anaphase onset, the sister chromatids then segregate, creating daughter cells with equal chromosome content. During meiosis, one round of DNA replication is followed by two rounds of chromosome segregation. First, the paired homologous chromosomes attach to microtubules emanating from opposite spindle poles and segregate in meiosis I. Then sister chromatid kinetochores attach to opposite spindle poles and separate in meiosis II, creating haploid meiotic products.

A crucial component of faithful chromosome segregation is the process of correcting erroneous kinetochore-microtubule attachments. For example, if both sister chromatid kinetochores are attached to microtubules emanating from the same spindle pole in mitosis or meiosis II, or if homologous chromosomes are attached to the same pole in meiosis I, the kinetochores are not under tension. A highly conserved kinase, Aurora B kinase (Ipl1 in budding yeast), releases tension-less attachments by phosphorylating kinetochore proteins to disrupt their interaction with microtubules (reviewed in (Cairo and Lacefield, 2020; Krenn and Musacchio, 2015)). The unattached kinetochores then recruit spindle checkpoint proteins, leading to the formation of the mitotic checkpoint complex and inhibition of the Anaphase Promoting Complex/Cyclosome (APC/C). Once bipolar attachments are made, protein phosphatase 1 (PP1) binds the kinetochore and dephosphorylates proteins to stabilize kinetochore-microtubule attachments and turn off the spindle checkpoint (Saurin, 2018). The APC/C then becomes active and targets proteins such as securin, an inhibitor of separase, for ubiquitination and subsequent proteasomal degradation. Separase is then free to cleave cohesin, allowing chromosome segregation in anaphase.

Localization of Aurora B to the inner centromere or kinetochore is important for its role in error correction. Aurora B kinase is part of the chromosomal passenger complex (CPC), which is required for both localization and Aurora B kinase activity (Cairo and Lacefield, 2020). Studies from mammalian cells lines, *Xenopus* egg extracts, budding yeast, and fission yeast have revealed two important pathways for Aurora B localization, the Bub1/Bub3 pathway and the haspin kinase pathway (Edgerton et al., 2016; Jeyaprakash et al., 2011; Kawashima et al., 2010; Kelly et al., 2010; Wang et al., 2010; Yamagishi et al., 2010). However, whether the haspin kinase pathway is functional for Ipl1^Aurora B^ inner centromere recruitment in budding yeast is currently debated (Campbell and Desai, 2013; Edgerton et al., 2016; Galli et al., 2020; Nespoli et al., 2006; Panigada et al., 2013). In addition, budding yeast have two additional pathways that control Ipl1^Aurora B^ localization, through the inner centromere protein Ndc10 and through the COMA complex (Cho and Harrison, 2011; Fischbock-Halwachs et al., 2019; Garcia-Rodriguez et al., 2019; Yoon and Carbon, 1999). Whether these four pathways contribute to Aurora B’s role in error correction during meiosis compared to mitosis has not been tested.

Each of the Aurora B recruitment pathways may bring a spatially distinct pool of Aurora B to the inner centromere or kinetochore (Figure 1A). In the Bub3/Bub1 recruitment pathway, Bub3 and Bub1 bind the kinetochore and Bub1 phosphorylates histone H2A (Baron et al., 2016; Kawashima et al., 2010; Kitajima et al., 2005; Kitajima et al., 2006; Liu et al., 2013a; Liu et al., 2015; Xu et al., 2009; Yamagishi et al., 2010). Shugoshin binds phosphorylated histone H2A and recruits the CPC to the inner centromere (Abad et al., 2022; Gassmann et al., 2004; Liu et al., 2013b; Sampath et al., 2004; Tsukahara et al., 2010). In the haspin kinase pathway, haspin kinase phosphorylates histone H3, which serves as a receptor for CPC binding (Dai et al., 2005; Edgerton et al., 2016; Jeyaprakash et al., 2011; Kelly et al., 2010; Wang et al., 2010; Yamagishi et al., 2010). In the budding yeast-specific Ndc10 pathway, the centromere-binding protein Ndc10 can interact with the CPC component Bir1 to bring the CPC to the inner centromere (Cho and Harrison, 2011; Yoon and Carbon, 1999). Finally, in the budding yeast-specific COMA complex (containing Ctf19, Okp1, Mcm21, and Ame1) pathway, Sli15 binds to Ctf19 to help localize Ipl1^Aurora B^ (Fischbock-Halwachs et al., 2019; Garcia-Rodriguez et al., 2019). Whether each of these kinetochore-localized pools of Aurora B contributes to error correction of kinetochore-microtubule attachment in meiosis is unknown.

**Figure 1.**
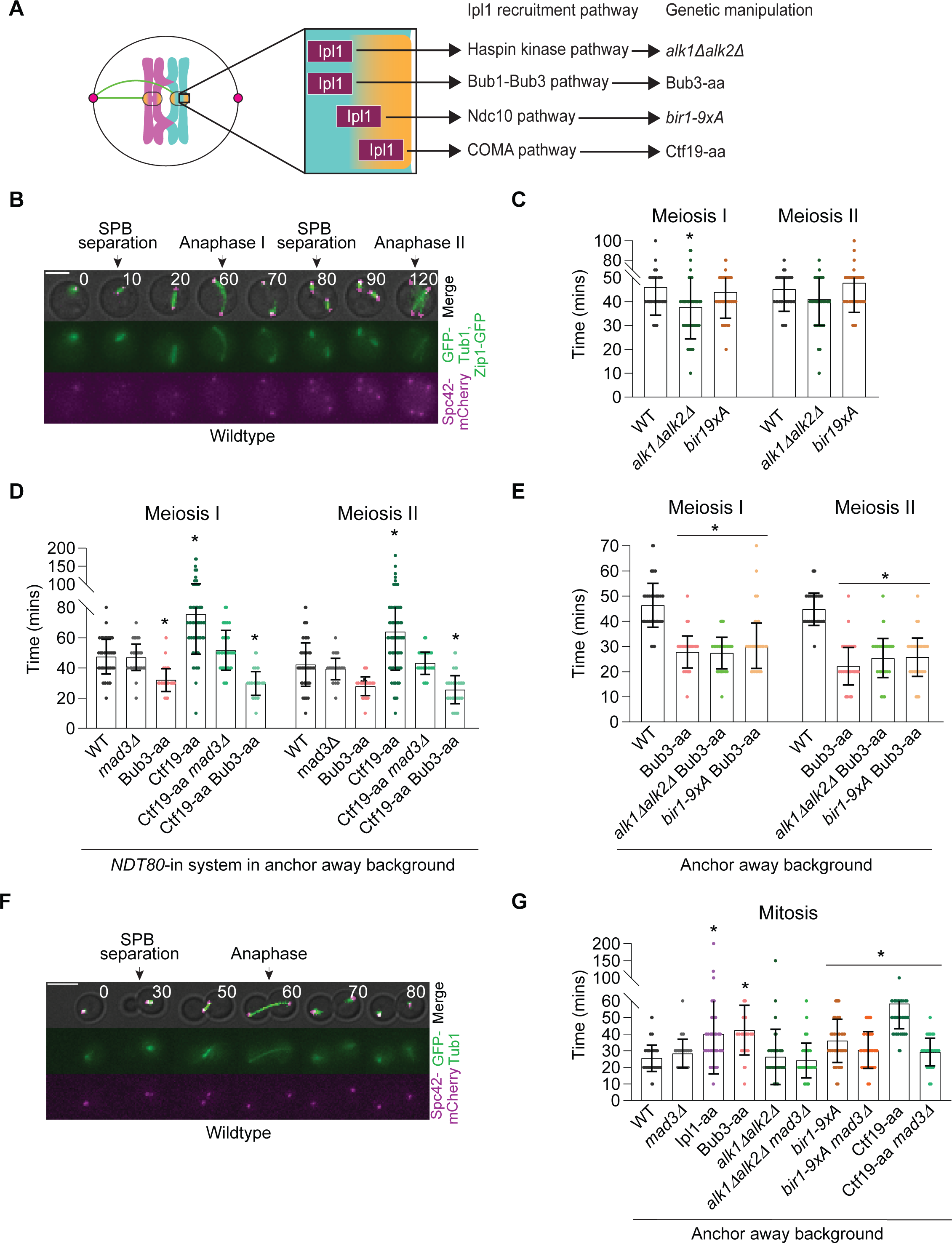
The Bub1/Bub3 pathway is the most important for setting the duration of mitosis and meiosis, however in opposing ways. (A) Cartoon indicating which genetic manipulation was used to disrupt each Ipl1 recruitment pathway. (B) Representative time-lapse images of a wildtype cell undergoing meiosis. Cell expresses *ZIP1-GFP*, *GFP-TUB1* and *SPC42-mCherry*. Time 0 marks prophase I. Numbers indicate time in minutes. Scale bar, 5μm. (C) Graph of the mean time from SPB separation to anaphase onset in meiosis I and meiosis II. * indicates a statistically significant difference compared to wildtype cells (at least 100 cells from two or more independent experiments per genotype were counted; P<0.05, Mann-Whitney test; error bars show standard deviation (SD)). (D) Graph of the mean time from SPB separation to anaphase onset in meiosis I and meiosis II. Strains contain the *NDT80*-in system (*P_GAL1,10_-NDT80, Gal4-ER*) to synchronize cells and the anchor away background (*TOR1-1*, *fpr1Δ*, *RPL13-2XFKBP12*). In all strains, except wildtype and *mad3Δ*, rapamycin was added 45 minutes after prophase I release to deplete Bub3 and/or Ctf19 from the nucleus. * indicates a statistically significant difference compared to wildtype cells (at least 100 cells from two or more independent experiments per genotype were counted; P<0.05, Mann-Whitney test; error bars show SD). aa, anchor away. (E) Graph of the mean time from SPB separation to anaphase onset in meiosis I and meiosis II. Strains contain the anchor away background (*TOR1-1*, *fpr1Δ*, *RPL13-2XFKBP12*). Rapamycin was added when cells were transferred into sporulation medium in strains containing Bub3-aa to deplete Bub3 from the nucleus. * indicates a statistically significant difference compared to wildtype cells (at least 100 cells from two or more independent experiments per genotype were counted; P<0.05, Mann-Whitney test; error bars show SD). aa, anchor away. (F) Representative time-lapse of a wildtype cell going through mitosis. Cell expresses *GFP-TUB1* and *SPC42-mCherry*. Numbers indicate time in minutes. Scale bar, 5μm. (G) Graph of the mean time from SPB separation to anaphase onset in mitosis. Strains have the anchor away background. In strains containing Ipl1-aa, Bub3-aa, and Ctf19-aa, rapamycin was added when diluting (1:20) an overnight culture, 40-45 minutes before imaging to deplete Ipl1, Bub3, and Ctf19 from the nucleus, respectively. * indicates a statistically significant difference compared to wildtype cells (at least 100 cells from two or more independent experiments per genotype were counted; P<0.05, Mann-Whitney test; error bars show SD). aa, anchor away.

Interestingly, the pools of Ipl1^Aurora B^ brought by the distinct recruitment pathways may have different or more important roles in meiosis compared to mitosis. In our previous study, we analyzed the consequence of loss of the Bub3/Bub1 recruitment pathway and found two significant differences between mitosis and meiosis (Cairo and Lacefield, 2020; Cairo et al., 2020). First, depletion of Bub3/Bub1 caused faster anaphase I and anaphase II onset, but delayed mitotic anaphase onset (Cairo et al., 2020; Yang et al., 2015). The premature anaphase onset in meiosis was not due to loss of spindle checkpoint activity; instead, low levels of kinetochore-localized Ipl1^Aurora B^ resulted in premature kinetochore-localization of PP1, causing the faster anaphase onset (Cairo et al., 2020). Second, disruption of the Bub3/Bub1 pathway caused massive chromosome mis-segregation in meiosis but not in mitosis, in which more chromosomes traveled to one spindle pole in both meiosis I and meiosis II. This “uneven” chromosome segregation is due to a failure in error correction of inappropriate kinetochore-microtubule attachments. In budding yeast meiosis, most chromosomes initially attach to the old spindle pole body (SPB) rather than the newly duplicated SPB. Ipl1^Aurora B^ is needed to release the improper attachments so that bipolar attachments can be made. In the absence of Bub3/Bub1, the levels of Ipl1^Aurora B^ was significantly reduced on meiotic kinetochores but not mitotic kinetochores.

In this study, we asked whether the other pools of Ipl1^Aurora B^, brought to the inner centromere and kinetochore through the four different recruitment pathways are also important for normal timing of anaphase onset and for error correction of kinetochore-microtubule attachments in meiosis and mitosis in budding yeast. By assessing mutants or depletion of each pathway, we find the Bub1/Bub3 pathway is the most important for setting the normal duration of meiosis and mitosis, but through different mechanisms. For error correction of kinetochore-microtubule attachments, the pools of Ipl1^Aurora B^ brought by Bub3/Bub1 and COMA recruitment pathways are individually important in meiosis, but are compensatory in mitosis. Finally, we studied the role of the haspin and Bub1 recruitment pathways in mammalian oogenesis and found that the two pathways have a compensatory role in error correction in meiosis I and meiosis II. Our results demonstrate that the pools of Aurora B at the kinetochore can contribute to different functions of Aurora B and the importance of each pool can differ between mitosis and meiosis and among organisms.

## Results

### The pool of Ipl1^Aurora B^ at the inner centromere recruited through Bub1/Bub3 is the most important pool for setting the duration of meiosis I and meiosis II

In our previous study, we found that depleting Bub3 caused decreased levels of inner centromere-localized Ipl1^Aurora B^, resulting in a faster anaphase I and anaphase II onset (Cairo et al., 2020). These results led us to question whether other pools of Ipl1^Aurora B^, recruited to the inner centromere or kinetochore through different recruitment pathways, could also regulate the timing of anaphase onset in meiosis. To this end, we made mutants to disrupt centromere and kinetochore localization of Ipl1^Aurora B^ through the haspin kinase, Ndc10, and COMA complex localization pathways (Figure 1A). The cells also expressed *GFP-TUB1,* which marks the spindle, Zip1-GFP, which marks the synaptonemal complex, and *SPC42-mCherry* which shows the SPBs (Figure 1B). Although Tub1 and Zip1 are both tagged with GFP, the proteins are temporally and morphologically distinct, in that the synaptonemal complex assembles and disassembles in prophase I, prior to spindle formation (Tsuchiya and Lacefield, 2013). We then measured the time from SPB separation to anaphase spindle elongation in meiosis I and meiosis II (Figure 1B).

First, to disrupt the inner centromere recruitment of Ipl1^Aurora B^ through the haspin kinases, we deleted both genes encoding haspin kinases, *ALK1* and *ALK2* (Higgins, 2003). The loss of haspin kinases caused only an approximately 10-minute decrease in the time of anaphase I onset, when compared to wildtype cells (Figure 1C). The duration of meiosis II was not significantly different from wildtype. Therefore, the haspin kinases have only a minor role in setting the duration of meiosis I.

To disrupt the inner kinetochore localization of Ipl1^Aurora B^ through Ndc10, we analyzed a *bir1-9xA* strain. In this strain, the CPC component Bir1 has nine CDK phosphorylation sites mutated to alanine (S383, S395, S552, S587, S667, T684, S688, T735, and T747), disrupting the interaction with Ndc10 (Widlund et al., 2006). The *bir1-9xA* mutant undergoes anaphase I and anaphase II onset with similar timings to wildtype cells (Figure 1C). These results suggest that the pool of Ipl1^Aurora B^ brought to the inner kinetochore by Ndc10 is not important for the normal duration of meiosis I and meiosis II.

Finally, we disrupted the COMA complex recruitment of Ipl1^Aurora B^ to the kinetochore by depleting Ctf19 from the nucleus in prometaphase I using the anchor-away technique (Haruki et al., 2008); these cells expressed Ctf19 tagged with FRB (FKBP12-rapamycin-binding domain), and the ribosomal protein Rpl13a tagged with FKBP12. Rapamycin addition leads to the stable interaction of FRB and FKBP12, which results in the removal of Ctf19 from the nucleus along with the ribosomal subunit (Haruki et al., 2008). Although Ctf19 is a non-essential kinetochore component in mitosis, the kinetochore is extensively reorganized during meiosis and Ctf19 has a crucial role in kinetochore re-assembly after cells exit prophase I (Borek et al., 2021; Hyland et al., 1999; Ortiz et al., 1999). To avoid altering kinetochore structure, we depleted Ctf19 from the nucleus after kinetochore re-assembly. We synchronized cells in prophase I using a *GAL-NDT80* strain background expressing GAL4 fused to the estrogen receptor (Gal4-ER). The cells were arrested in prophase I, due to a lack of the Ndt80 transcription factor, and then were released from the arrest with the addition of β-estradiol (Benjamin et al., 2003; Carlile and Amon, 2008; Xu et al., 1995). By adding rapamycin 45-minutes after β-estradiol addition, we depleted Ctf19-FRB from the nucleus just before prometaphase I. We then monitored cells for the timing of anaphase I and anaphase II onset. The cells were strongly delayed in anaphase I and anaphase II onset when compared to the wildtype control cells that have Rpl13-FKB12 tagged, but without Ctf19 tagged (Figure 1D).

We considered that the loss of Ctf19 could cause defects in kinetochore-microtubule attachments that resulted in spindle checkpoint activation. We therefore deleted the spindle checkpoint protein *MAD3* in the Ctf19-FRB strain and found that cells underwent anaphase I and anaphase II with timings similar to wildtype (Figure 1D). This contrasted with Bub3-depleted cells, which also lacks spindle checkpoint activity, but displayed faster anaphase onset due to the loss of Ipl1^Aurora B^ kinetochore localization (Figure 1D) (Cairo et al., 2020). Therefore, the delay in anaphase I and anaphase II onset in Ctf19-depleted cells was dependent on the spindle checkpoint. We conclude that the loss of the COMA-dependent pool of Ipl1^Aurora B^ at the kinetochore does not result in a faster anaphase I or anaphase II onset, even with the disruption of the spindle checkpoint.

To determine if there was an additive affect from losing two Ipl1^Aurora B^ pools, we depleted Bub3 and the components of the other pathways. We found that the timings of anaphase I and anaphase II onset with loss of Bub3 and haspin kinase pathways, Bub3 and Ndc10 pathways, as well as Bub3 and COMA pathways were similar to the timings of Bub3-depleted cells (Figure 1C-E). These results suggest that loss of the other pathways does not cause an enhanced defect in combination with loss of the Bub3/Bub1 recruitment pathway. Overall, we conclude that the Ipl1^Aurora B^ pool recruited to the inner centromere through the Bub3/Bub1 pathway is the most important for the normal timing of anaphase I and anaphase II onset.

In mitosis, we previously showed that loss of Bub1/Bub3 resulted in slower anaphase onset instead of the faster anaphase onset displayed in meiosis (Cairo et al., 2020; Yang et al., 2015). We next asked if loss of Ipl1^Aurora B^ also delayed anaphase onset. Using anchor-away, we depleted Ipl1^Aurora B^ from the nucleus and measured the time from SPB separation to spindle elongation at anaphase onset. Compared to the wildtype control strain, nuclear depletion of Ipl1^Aurora B^ delayed anaphase onset, similar to Bub3 nuclear depletion (Figure 1F-G).

Therefore, we next asked if loss of other Ipl1^Aurora B^ recruitment pathways also elongated mitosis. The *alk1Δ alk2Δ* cells had a similar time of anaphase onset as wildtype cells (Figure 1G). In contrast, *bir1-9xA* mutants and Ctf19-depleted cells had a delayed anaphase onset, similar to the time of anaphase onset in Bub3 and Ipl1-depleted cells (Figure 1G). However, the delay in *bir1-9xA* mutants and Ctf19-depleted cells was at least partially dependent on the spindle checkpoint; deletion of *MAD3* decreased the time of anaphase onset (Figure 1G).

We conclude that Ipl1^Aurora B^ is also important for setting the duration of mitosis, likely through the pool localized to the kinetochore by Bub3/Bub1. However, loss of Ipl1^Aurora B^ leads to a slower anaphase onset in mitosis and a faster anaphase onset in meiosis, suggesting that the mechanism of regulation may be different between the two types of cell cycles.

### PP1 kinetochore localization and the phospho-regulation of Cdc20 sets the duration of anaphase onset in meiosis but not mitosis

We previously showed that faster anaphase I and anaphase II onset in Bub3 depleted cells is due to the loss of kinetochore localized Ipl1^Aurora B^, which causes premature kinetochore localization of PP1 (Cairo et al., 2020).Our next goal was to identify the substrate of PP1 that leads to premature anaphase onset in meiosis. Studies in mitosis in *C. elegans* and mammalian cell lines showed that kinetochore-localized PP1 can dephosphorylate Cdc20 so that it can bind and activate the APC/C to promote anaphase onset (Bancroft et al., 2020; Kim et al., 2017). In budding yeast, Cdc20 phosphorylation and dephosphorylation has not been shown to regulate the timing of anaphase onset in mitosis. However, a recent genome-wide phosphoproteomics study identified several phosphorylation sites in budding yeast Cdc20 (Lanz et al., 2021). Therefore, we asked if Cdc20 phosphorylation and dephosphorylation regulates the timing of anaphase onset in mitosis and meiosis.

We investigated the role of three phosphorylation sites identified in the N-terminus of Cdc20 by the phospho-proteomics study, S127, S88, and S89 (Lanz et al., 2021). We chose to analyze S127 because it is adjacent to a conserved motif that aligns with a CDK phosphorylation site important for regulating the timing of anaphase onset in human cells and *C. elegans* mitosis (Bancroft et al., 2020; Kim et al., 2017) (Figure 2A). S88 and S89 are in a basic patch consensus motif that has similarities with other known PP1 binding motifs, although the sequence does not contain the exact PP1 binding motif (Figure 2A) (Heroes et al., 2013; Smith et al., 2019). We hypothesize that Cdc20 dephosphorylation at S127 by PP1 promotes anaphase onset. This hypothesis leads to the prediction of premature anaphase onset if Cdc20 could not be phosphorylated at S127; conversely, mutating S88, S89 would delay anaphase onset due to decreased PP1 binding.

**Figure 2.**
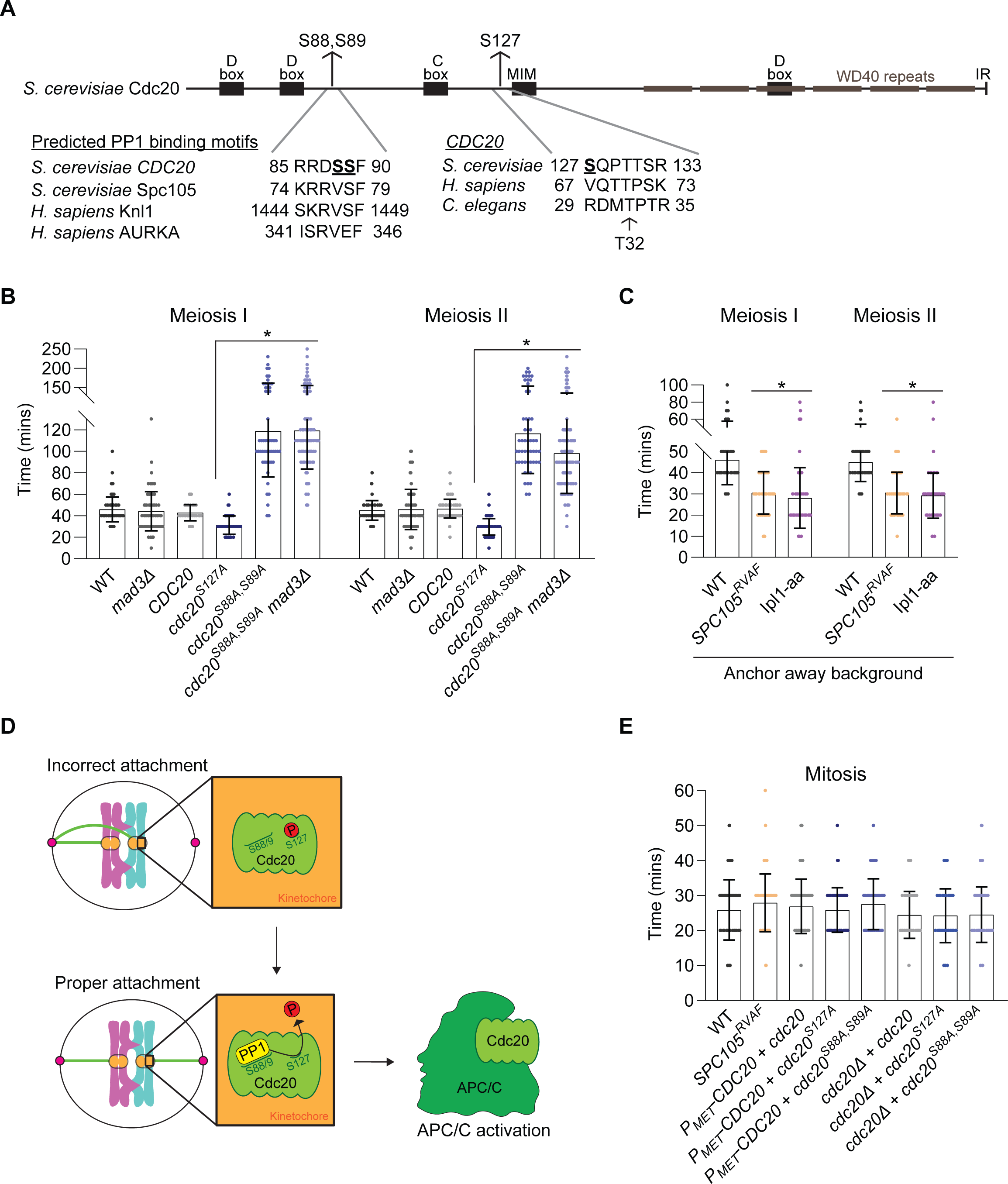
PP1 kinetochore localization and Cdc20 dephosphorylation sets the timing of anaphase onset in meiosis but not mitosis. (A) Diagram of Cdc20 domains highlighting the N-terminal phosphorylation sites S88, S89, and S127, predicted PP1 binding motifs in *S. cerevisiae* and *H. sapiens*, and *S. cerevisiae CDC20* alignment with *H. sapiens and C. elegans CDC20*. (B) Graph of the mean time from SPB separation to anaphase onset in meiosis I and meiosis II. All strains, except wildtype and *mad3Δ*, have the endogenous *CDC20* under the mitosis-specific *CLB2* promoter and express an integrated *CDC20*, *cdc20^S127A^* or *cdc20^S88A,S89A^* under the *CDC20* promoter. * indicates a statistically significant difference compared to wildtype cells (at least 100 cells from two or more independent experiments per genotype were counted; P<0.05, Mann-Whitney test; error bars show SD). (C) Graph of the mean time from SPB separation to anaphase onset in meiosis I and meiosis II. All strains contain the anchor away genetic background (*tor1-1*, *fpr1Δ*, *RPL13-2XFKBP12*). In *SPC105^RVAF^* cells, rapamycin was added 7 hours after resuspension into sporulation media to allow enough *SPC105^RVAF^* protein to be produced before anchoring away the FRB-tagged Spc105. In Ipl1-aa cells, rapamycin was added when cells were resuspended in sporulation medium to deplete Ipl1 from the nucleus. * indicates a statistically significant difference compared to wildtype cells (at least 100 cells from two or more independent experiments per genotype were counted; P<0.05, Mann-Whitney test; error bars show SD). aa, anchor away. (D) Cartoon depicting proposed meiotic model. When bipolar attachments are established, PP1 binds the kinetochore and dephosphorylates kinetochore-localized Cdc20 for APC/C activation. (E) Graph of the mean time from SPB separation to mitotic anaphase onset. Strains containing *P_MET_CDC20* have the endogenous *CDC20* under the *MET3-*repressible promoter and express an integrated *CDC20*, *cdc20^S127A^* or *cdc20^S88AS89A^* under the *CDC20* promoter. In strains containing *cdc20Δ*, the endogenous CDC20 is deleted and integrated *cdc20*, *cdc20^S127A^* or *cdc20^S88AS89A^* is under the endogenous promoter. At least 100 cells from two independent experiments per genotype were counted; P<0.05, Mann-Whitney test; error bars show SD.

To test this hypothesis, we expressed a mutant version of *CDC20* with serine 127 or serines 88 and 89 changed to alanine (*cdc20^S127A^* or *cdc20^S88A,S89A^*). These strains had the wildtype copy of *CDC20* under the mitosis-specific *CLB2* promoter to prevent expression of wildtype *CDC20* in meiosis (Carlile and Amon, 2008; Dahmann and Futcher, 1995; MacKenzie and Lacefield, 2020). As a control, we integrated a wildtype copy of *CDC20* into the strain with *CLB2* promoted *CDC20*. We then measured the time from SPB separation to anaphase onset. We found that *cdc20^S127A^* cells underwent anaphase I and anaphase II onset approximately 12 and 17 minutes faster than the *CDC20* control cells, respectively (Figure 2B). The faster metaphase I and metaphase II did not disrupt chromosome segregation in that the spores showed similar viability as wildtype cells (Figure S1A).

We next compared the timing of anaphase onset in cells depleted of Ipl1^Aurora B^ and in *SPC105^RVAF^* cells. With Ipl1^Aurora B^ depletion, PP1 likely localizes to the kinetochore prematurely (Cairo et al., 2020; Rosenberg et al., 2011). Similarly, the mutation in the RVSF motif to RVAF prevents inhibitory phosphorylation of PP1s binding site by Ipl1^Aurora B^, which should allow PP1 to localize prematurely (Figure 2C) (Hendrickx et al., 2009; Liu et al., 2010; Rosenberg et al., 2011). We depleted Ipl1^Aurora B^ from the nucleus using anchor-away (Haruki et al., 2008).

Similarly, anchor away was also used in *SPC105^RVAF^* cells to deplete the wildtype Spc105 while expressing *SPC105^RVAF^* from the meiosis-specific *REC8* promoter. Strikingly, the timing of anaphase I and anaphase II onset of cells expressing *cdc20^S127A^* was similar to that of Ipl1^Aurora B^ depleted and *SPC105^RVAF^* cells (Figure 2B-C).

*cdc20^S88A, S89A^* cells were strongly delayed, with approximately 120 minutes to anaphase I onset and 115 minutes to anaphase II onset when compared to wildtype (Figure 2B). To test if this delay was spindle checkpoint dependent, we deleted *MAD3* in *cdc20^S88A,S89A^* cells and observed an approximately 120 minute and 100 minute delay in anaphase I and anaphase II onset, respectively (Figure 2B). Additionally, we noticed that approximately 3% of *cdc20^S88A, S89A^ mad3Δ* cells arrested after spindle breakdown in meiosis I and 6% arrested in metaphase II. These results show that the delay observed in *cdc20^S88A,S89A^* cells is not spindle checkpoint dependent.

Taken together, our results support a model that PP1 binds Cdc20 at S88 and S89 and dephosphorylates S127 to promote anaphase I and anaphase II onset (Figure 2D). However, we cannot exclude the possibility that PP1 is acting indirectly on Cdc20.

We next analyzed the timing of mitotic anaphase onset in *cdc20^S127A^* and *cdc20^S88A,S89A^* cells expressing *GFP-TUB1* and *SPC42-mCherry* (Figure 2E). To this end, we placed the endogenous *CDC20* under the control of the *MET3-*repressible promoter and integrated into the genome *cdc20^S127A^* or *cdc20^S88A,S89A^* under the *CDC20* promoter. In this strain, the endogenous *CDC20* should not be expressed in the presence of methionine (Uhlmann et al., 2000). As a control, we integrated a plasmid with *CDC20* in the same *MET3*-promoted *CDC20* strain. We released cells from an α-factor arrest in medium containing methionine and monitored the cells for the timing of anaphase onset. The cells expressing *cdc20^S127A^* and *cdc20^S88A,S89A^* underwent anaphase onset with similar timings to cells expressing *CDC20* (Figure 2E).

The difference between meiosis and mitosis in the timing of anaphase onset in cells expressing *cdc20^S127A^* was surprising. To ensure that the normal anaphase onset timings in the *MET3-*repressible *CDC20* strain was not due to a low level of *CDC20* expression, we deleted the endogenous copy of *CDC20* in the strains expressing *cdc20^S127A^* or *cdc20^S88A,S89A^.* We found that the time of SPB separation to anaphase onset in *cdc20^S127A^* and *cdc20^S88A,S89A^* cells was similar to those of cells expressing wildtype *CDC20,* which further suggested that S127 phosphorylation and dephosphorylation did not alter the timings of mitosis (Figure 2E). Our results support the model that phosphorylation and dephosphorylation of Cdc20 at S127 is important for setting the normal duration of meiosis but not mitosis.

### The pools of Ipl1^Aurora B^ at the inner centromere and kinetochore recruited through Bub1/Bub3 pathway and the COMA pathway are important for error correction of kinetochore-microtubule attachments in meiosis I and meiosis II

Our previous study showed that loss of Ipl1^Aurora B^ kinetochore localization from the Bub1/Bub3 pathway caused a failure in error correction of kinetochore-microtubule attachments. We asked if the other pools of Ipl1^Aurora B^ at the centromere and kinetochore are also important for error correction (Cairo et al., 2020). To monitor chromosome segregation, we tagged the histone protein Htb2 with mCherry (Htb2-mCherry). Although we cannot see individual chromosomes, we can monitor bulk chromatin separation (Figure 3A-C). In budding yeast, most initial attachments are made to the older SPB that emanates more microtubules, and the error correction kinase Ipl1^Aurora B^ is required for releasing those initial attachments to make bipolar attachments (Meyer et al., 2013; Tanaka et al., 2002). In wildtype cells, two chromatin masses of equal size are present after meiosis I and four chromatin masses of equal size are present after MII (Figure 3B, D-E). As we previously observed, with depletion of Ipl1^Aurora B^ or Bub3, unequal chromatin mass sizes are present in both meiosis I and meiosis II, consistent with a failure in correction of the initial improper kinetochore-microtubule attachments at the old SPB (Figure 3D, F) (Cairo et al., 2020).

**Figure 3.**
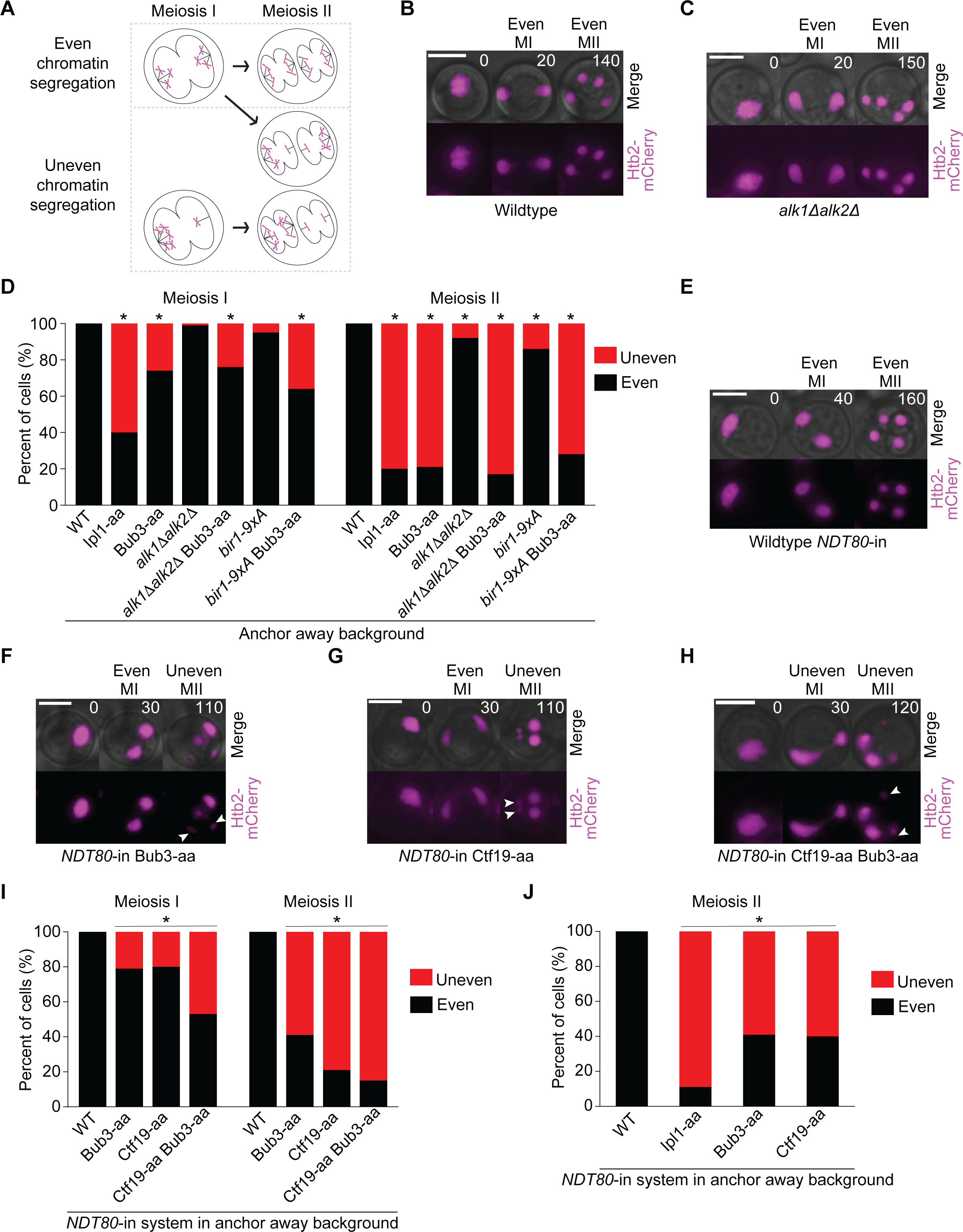
The pools of Ipl1^Aurora B^ at the inner centromere and kinetochore recruited through Bub1/Bub3 pathway and the Ctf19 pathway are important for error correction of kinetochore-microtubule attachments in meiosis I and meiosis II. (A) Schematic of even and uneven chromatin segregation in meiosis I and meiosis II. (B, C) Representative time-lapse of wildtype (B) and *alk1Δalk2Δ* (C) cells displaying even chromatin segregation in meiosis I and meiosis II. Cells express Htb2-mCherry to visualize chromatin. Numbers indicate time in minutes. Scale bars, 5μm. (D) Percent of cells that display even and uneven chromatin segregation in meiosis I and meiosis II. All strains contain the anchor away genetic background (*TOR1-1*, *fpr1Δ*, *RPL13-2XFKBP12*). Rapamycin was added to Bub3-aa and Ipl1-aa cells upon resuspension in sporulation medium to deplete Bub3 and Ipl1 from the nucleus, respectively. * indicates a statistically significant difference compared with the wildtype anchor away strain (at least 100 cells from two or more independent experiments per genotype were counted; P < 0.05; two-tailed Fisher’s exact test). aa, anchor away. (E-H) Representative time-lapse of wildtype (E), Bub3-aa (F), Ctf19-aa (G), and Ctf19-aa Bub3-aa (H) cells containing the *NDT80*-in system displaying either even or uneven chromatin segregation events in meiosis I and II. Cells express Htb2-mCherry. Numbers indicate time in minutes. Scale bars, 5μm. aa, anchor away. (I) Percent of cells that display even and uneven chromatin segregation in meiosis I and meiosis II, containing the *NDT80*-in system (*P_GAL1,10_-NDT80, Gal4-ER*) and the anchor away genetic background (*tor1-1*, *fpr1Δ*, *RPL13-2XFKBP12*). In all strains, β-estradiol was added 12 hours after cells were resuspended in sporulation media. Rapamycin was added to strains containing Bub3-aa and Ctf19-aa 45 minutes after β-estradiol addition to deplete Bub3 and Ctf19 from the nucleus, respectively. * indicates a statistically significant difference compared with the wildtype anchor away strain (at least 100 cells from two or more independent experiments per genotype were counted; P < 0.05; two-tailed Fisher’s exact test). aa, anchor away. (J) Percent of cells that display even and uneven chromatin segregation in meiosis II, containing the *NDT80*-in system (*P_GAL1,10_-NDT80, Gal4-ER*) and the anchor away genetic background (*TOR1-1*, *fpr1Δ*, *RPL13-2XFKBP12*). β-estradiol was added to *NDT80*-in cells 12 hours after suspension in sporulation media. Rapamycin was added 3 hours after β-estradiol addition to Ipl1-aa, Bub3-aa, and Ctf19-aa cells, as cells were completing meiosis I to deplete Ipl1, Bub3, and Ctf19 from the nucleus, respectively. * indicates a statistically significant difference compared with the wildtype anchor away strain (at least 100 cells from two or more independent experiments per genotype were counted; P < 0.05; two-tailed Fisher’s exact test). aa, anchor away.

We next asked if the loss of other kinetochore pools of Ipl1^Aurora B^ also caused differences in bulk chromatin segregation. We first monitored loss of the haspin and Ndc10 pathways with *alk1Δ alk2Δ* and *bir1-9xA* cells, respectively. We observed a small, but statistically significant, fraction of cells with uneven chromatin masses after meiosis II in both mutants (Figure 3D). Loss of the haspin and Ndc10 pathways, combined with Bub3-depletion displayed a similar phenotype to Bub3 depleted cells (Figure 3D).

To avoid disrupting kinetochore assembly, we depleted Ctf19-FRB from the nucleus 45 minutes after the release from a prophase I arrest. Even in this case, 17% of cells showed additional Htb2-mCherry DNA masses after meiosis II, suggesting some kinetochore defects (Figure S2A-B). To ask if kinetochore re-assembly was disrupted with Ctf19 depletion, we monitored the outer kinetochore component Ndc80-GFP. We measured the fluorescence intensity of Ndc80-GFP foci and found similar levels in wildtype cells and in cells in which Ctf19 was anchored away in prometaphase I (Figure S3A-B). These results suggest that with Ctf19 depletion, the kinetochores could re-assemble and recruit Ndc80. We decided to continue with the analysis, only scoring cells that did not have additional DNA masses. Although these cells may have somewhat of a kinetochore disruption, we would not expect stabilized attachments in which the majority of chromosomes segregate to one pole unless Ctf19 was important for localizing the pool of Ipl1^Aurora B^ needed for error correction.

We found that depletion of Ctf19 resulted in approximately 20% of cells with uneven chromatin masses in meiosis I and 80% of cells with uneven chromatin masses in meiosis II, similar to Bub3-depleted cells (Figure 3E-I). The double depletion of both Bub3 and Ctf19 resulted in an even higher percent of cells with uneven chromosome segregation in meiosis I and meiosis II (Figure 3I). These results suggest that the pools of Ipl1^Aurora B^ recruited by both Bub3 and Ctf19 are important for error correction of kinetochore-microtubule attachments to allow proper chromosome segregation.

Our results showed that depletion of Ctf19 in prometaphase I caused a severe chromosome segregation defect in meiosis I and, especially, meiosis II. However, because addition of rapamycin in prometaphase I results in depletion of Ctf19 in both meiosis I and meiosis II, we asked if depletion after meiosis I, but just before meiosis II, also resulted in a high percent of chromosome mis-segregation in meiosis II (Cairo et al., 2022; Cairo et al., 2020) (Figure 3J). We synchronized cells in prophase I and added rapamycin 3 hours after the cells were released, just as they were entering meiosis II. 61% of cells displayed massive chromatin mis-segregation upon depletion of Ctf19, similar to the cells with a depletion of Bub3 (Figure 3J). These results suggest that each individual pool of Ipl1^Aurora B^ brought to the kinetochore by Ctf19 or the inner centromere by Bub3 is important for correction of kinetochore-microtubule attachments, especially in meiosis II.

### The pools of Ipl1^Aurora B^ at the inner centromere and kinetochore recruited through Bub1/Bub3 and the Ctf19 pathways are compensatory for error correction of kinetochore-microtubule attachments in mitosis

We next tested the role of the different pools of Ipl1^Aurora B^ at the kinetochore during mitosis. We monitored Htb2-mCherry cells and measured the DNA mass in the mother and the bud (Figure 4A-D). We found that there were no major defects in chromatin segregation in the individual single mutants that disrupt one pool of Ipl1^Aurora B^ kinetochore localization (Figure 4E). However, depletion of both Ctf19 and Bub3 resulted in a strong defect in chromatin segregation, with 47% of cells with uneven chromatin masses between the mother and the bud, which agrees with previous findings in the literature (Garcia-Rodriguez et al., 2019). Loss of the haspin and Ndc10 pathways, combined with Bub3-depletion did not display an enhanced defect (Figure 4E). We conclude that the Ipl1^Aurora B^ pool brought to the inner centromere and kinetochore through the Bub3 and Ctf19 pathways are important for error correction, but are compensatory in their roles.

**Figure 4.**
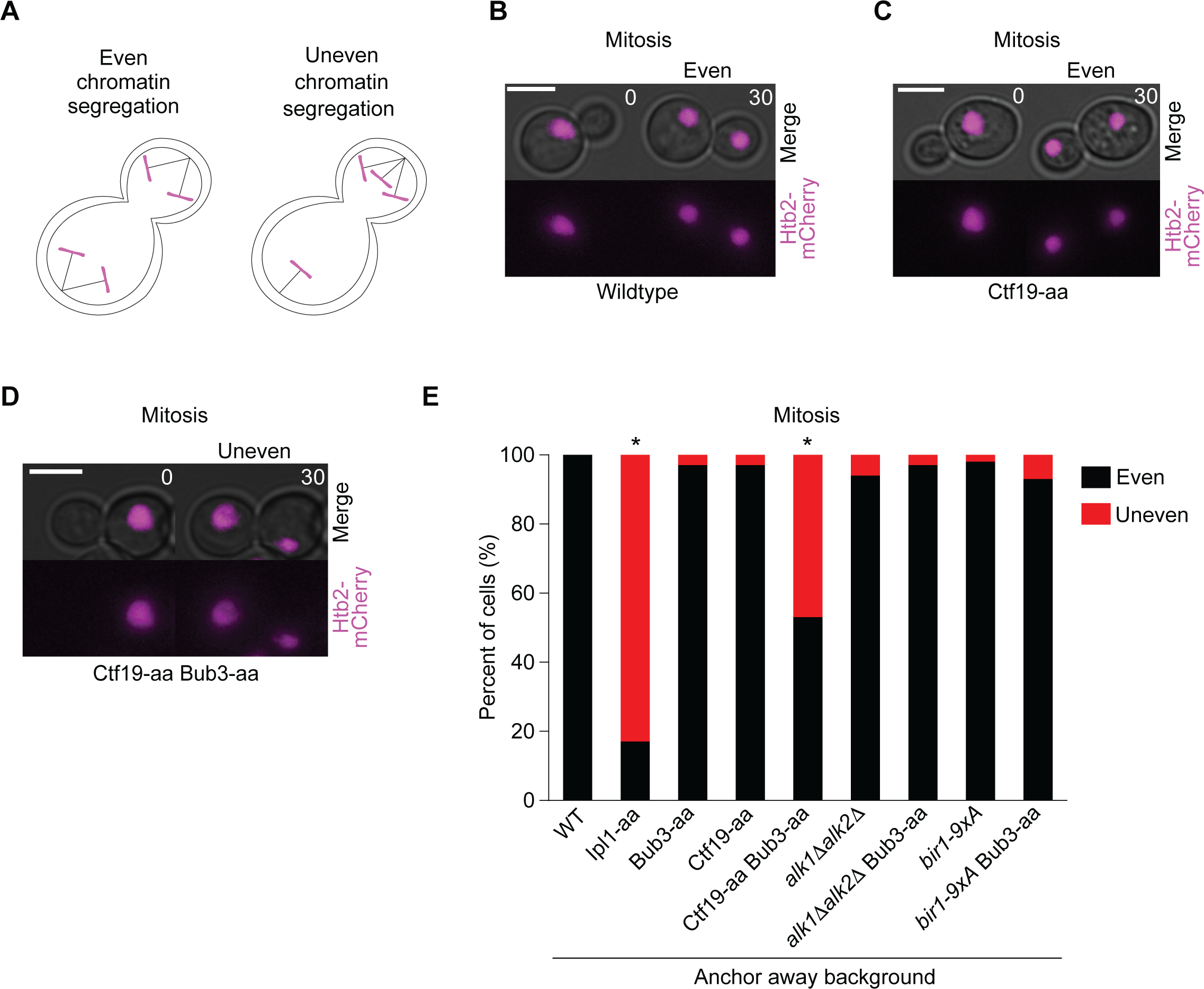
The pools of Ipl1^Aurora B^ at the inner centromere and kinetochore recruited through Bub1/Bub3 and the Ctf19 pathways are compensatory for error correction of kinetochore-microtubule attachments in mitosis. (A) Schematic of even and uneven chromatin segregation in mitosis. (B-D) Representative time-lapse of wildtype (B), Ctf19-aa (C), and Ctf19-aa Bub3-aa (D) cells expressing Htb2-mCherry and displaying even or uneven chromatin segregation. Numbers indicate time in minutes. Scale bars, 5μm. aa, anchor away. (E) Percent of cells displaying even and uneven chromatin segregation in mitosis. All strains contain the anchor away genetic background (*TOR1-1*, *fpr1Δ*, *RPL13-2XFKBP12*). Rapamycin was added to strains containing Ipl1-aa, Bub3-aa, and Ctf19-aa 30 minutes before imaging to deplete Ipl1, Bub3, and Ctf19 from the nucleus, respectively. * indicates a statistically significant difference compared with the wildtype anchor away cells (at least 100 cells from two or more independent experiments per genotype were counted; P < 0.05, two-tailed Fisher’s exact test). aa, anchor away. aa, anchor away.

### The pools of AURKC at the inner centromere recruited through Bub1 and haspin kinases are compensatory for error correction of kinetochore-microtubule attachments

We next wanted to compare our findings in budding yeast to a mammalian system. In mammalian cell lines, the two known Aurora B recruitment pathways are through haspin kinase phosphorylation of histone H3 and through Bub1 kinase phosphorylation of H2A. Inhibition of both pathways in the cell lines caused more severe defects in chromosome alignment and segregation than inhibition of a single pathway (Hadders et al., 2020; Liang et al., 2020). However, less is known about whether they have compensatory functions during mammalian oogenesis.

Past studies have shown that haspin kinase is important for normal spindle assembly and chromosome segregation (Nguyen et al., 2014). Inhibition of haspin kinases specifically in prometaphase I, after the spindle had formed, resulted in a loss of phosphorylation of histone H3 at threonine 3 (H3T3), and an increase in chromosome mis-segregation (Quartuccio et al., 2017). In contrast, analysis of oocytes with a deletion of the Bub1 kinase domain (*Bub1KD*) did not show a strong defect in chromosome mis-segregation. However, inhibition of both recruitment pathways has not been analyzed.

To this end, we chemically inhibited both haspin kinase and Bub1 kinase activity in oocytes matured *in vitro* (Figure 5A). For inhibition of haspin kinase, we added 0.5μM 5-iodotubercidin (5-Itu), which abrogates haspin kinase activity and therefore phosphorylation of H3T3 in oocytes (Nguyen et al., 2014). To inhibit Bub1 kinase activity, we added the compound BAY-1816032, which was previously characterized as a highly selective Bub1 kinase inhibitor in human cell lines (Siemeister et al., 2019). To determine the correct concentration of BAY-1816032 for Bub1 inhibition in mouse oocytes, we assessed the levels of H2AT120 phosphorylation after adding different concentrations of the drug. With a phosphorylation-specific antibody, we found that the levels of kinetochore-localized H2AT120 phosphorylation were significantly reduced at 8μM (Figure S4A-B).

**Figure 5.**
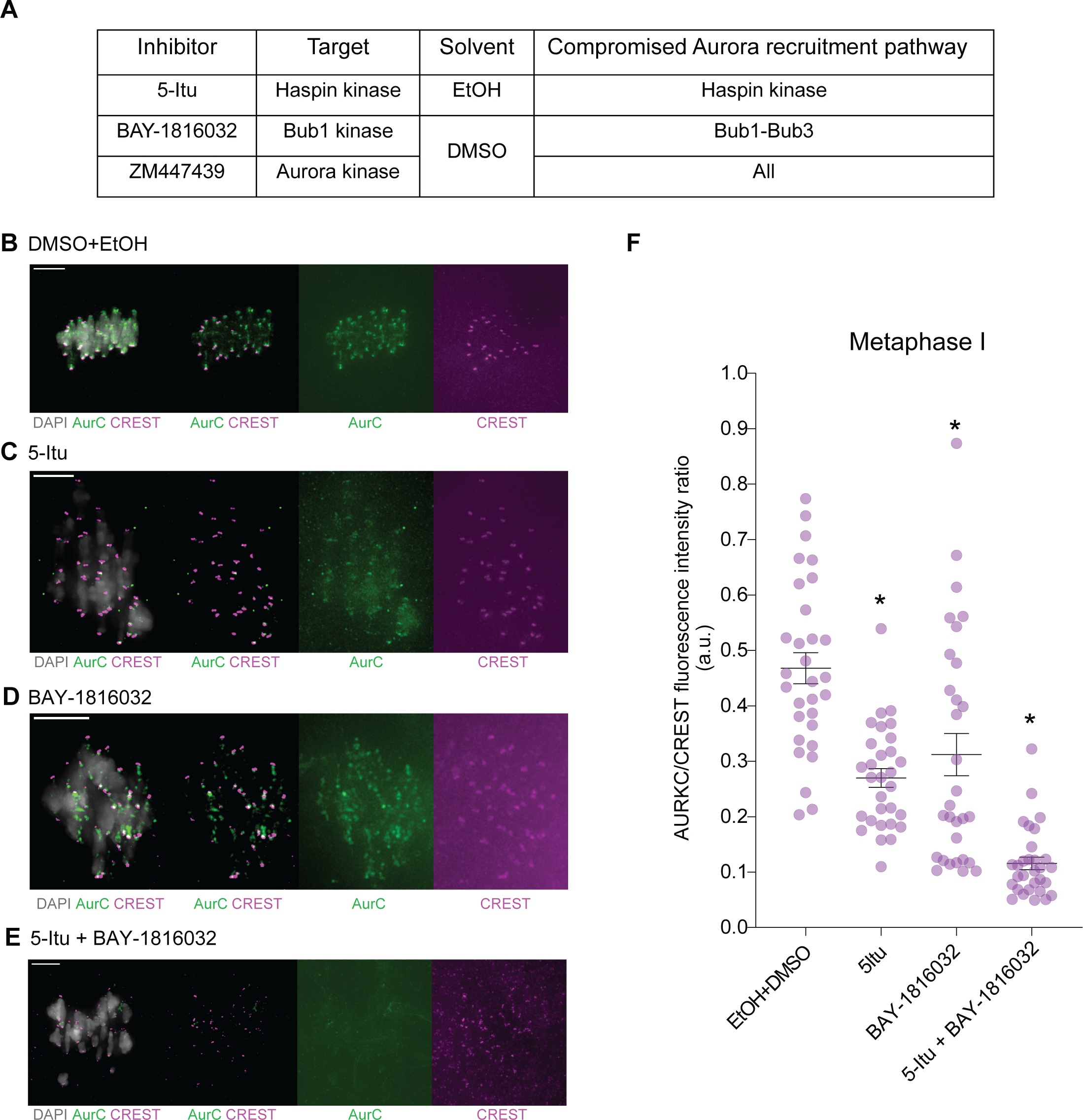
Bub1 and haspin kinases are compensatory for recruiting AURKC at the kinetochore. (A) Table depicting which inhibitor and corresponding solvent was used to disrupt each Aurora recruitment pathway in mouse oocytes. (B-E) Representative images of oocytes that were matured *in vitro* for 8 hours from prophase I release. Oocytes were treated with the indicated drugs or solvents and immunostained for DNA (DAPI, gray), Aurora kinase C (α-tubulin, green), and kinetochores (CREST, magenta). DMSO and BAY-1816032 were added in prophase I upon oocyte transfer into the maturation media; ethanol and 5-Itu were added 5 hours after prophase I release. Scale bars, 10μm. (F) Graph indicating Aurora kinase C fluorescence intensity relative to kinetochores (CREST) staining. Each point plotted represents the average of at least 20 kinetochores in a single oocyte. 30 metaphase I oocytes were used for each condition. * indicates a statistically significant difference compared with the wildtype oocytes (P < 0.05, two-tailed Fisher’s exact test).

We next measured the intensity of AURKC at the inner centromere in the presence of the inhibitors or the solvent controls, DMSO and ethanol, which were used to dissolve the inhibitors (Figure 5A). We found that with addition of 5-Itu or BAY-1816032, the AURKC intensity that overlapped with the CREST kinetochore focus was significantly reduced in comparison to the solvent controls (Figure 5B-F). We note that 5-Itu addition also disrupts AURKC localization to the interchromatid axis, as previously shown (Quartuccio et al., 2017). Addition of both 5-Itu and BAY-1816032 further decreases the amount of AURKC that co-localizes with CREST. These results support the model that both haspin and Bub1 kinase recruits distinct pools of AURKC to the inner centromere.

To analyze whether error correction is affected by inhibition of the kinases, we assayed chromosome alignment at metaphase I and metaphase II. We predicted that if error correction activity was reduced, there would be an increase in misaligned chromosomes. We measured the spindle length from the spindle midzone to the pole. We then scored oocytes as having misaligned chromosomes if they had one or more chromosomes that were one third the length of the spindle or greater away from the midzone (Figure 6A-B). We also included oocytes with one or more unattached chromosomes, in which chromosomes were not on the spindle. To assess meiosis I, we added BAY-1816032 as we released the oocytes from a prophase I arrest in culture. Because haspin kinase is important for meiotic resumption, spindle assembly and chromosome condensation, 5-Itu was added to oocytes 5hrs after release from prophase I, when the oocytes were in late prometaphase I (Nguyen et al., 2014; Quartuccio et al., 2017). We then scored metaphase I chromosome alignment 3 hours later. To assess chromosome alignment in metaphase II, we added the drugs after 11 hours, a timepoint in which the oocytes had completed anaphase I. We then scored the metaphase II oocytes 4 hours later. As controls, we added ethanol and DMSO to the oocytes, which were the solvents used to dissolve 5-Itu and BAY-1816032, respectively.

**Figure 6.**
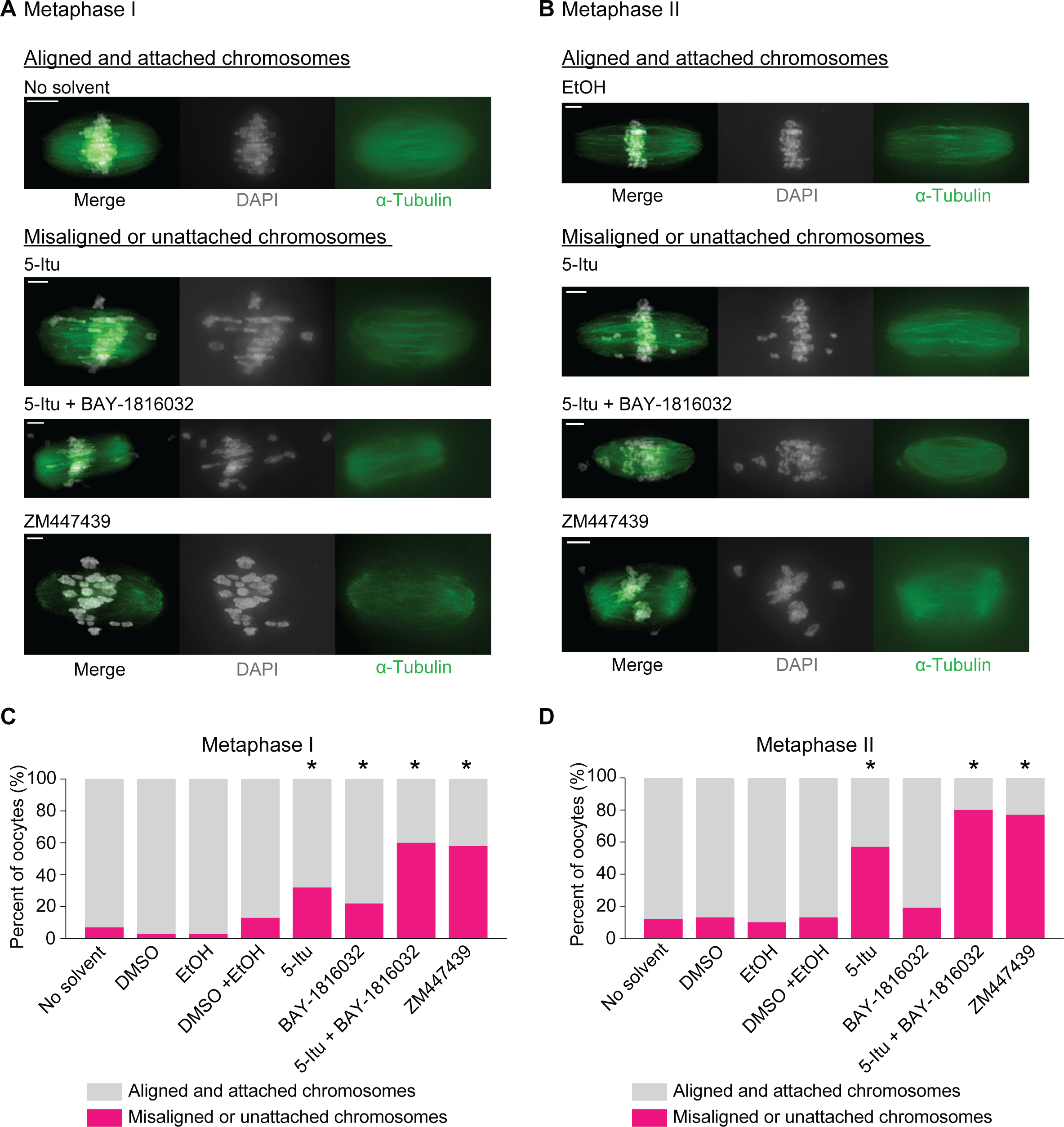
Bub1 and haspin kinases are compensatory for recruiting AURKC for error correction of kinetochore-microtubule attachments. (A) Representative images of oocytes that were matured *in vitro* for 8 hours from prophase I release. Oocytes were treated with the indicated drugs and immunostained for DNA (DAPI, gray) and microtubules (α-tubulin, green). DMSO, BAY-1816032, and ZM447439 were added in prophase I upon oocyte transfer into the maturation media; ethanol and 5-Itu were added 5 hours after prophase I release. Scale bars, 10μm. (B) Representative images of oocytes that were matured *in vitro* for 15 hours from prophase I release. Oocytes were treated with the indicated drugs and immunostained for DNA (DAPI, gray) and microtubules (α-tubulin, green). All drugs and solvents were added 11 hours after prophase I release. Scale bars, 10μm. (C) Percent of metaphase I oocytes with correctly aligned and attached or misaligned and unattached chromosomes. * indicates a statistically significant difference compared with the wildtype oocytes (n = 30-47 oocytes per condition; P < 0.05, two-tailed Fisher’s exact test). (D) Percent of metaphase II oocytes with correctly aligned and attached or misaligned and unattached chromosomes in metaphase I with the indicated drugs and solvents. * indicates a statistically significant difference compared with the wildtype oocytes (n = 30-47 oocytes per condition; P < 0.05, two-tailed Fisher’s exact test).

As expected from past studies, we found that the no solvent, ethanol, and DMSO control oocytes displayed 3-12% misaligned chromosomes in metaphase I and metaphase II (Nguyen et al., 2014; Quartuccio et al., 2017; Shuda et al., 2009) (Figure 6C-D). Addition of haspin kinase inhibitor resulted in 32% of oocytes with misaligned chromosomes in metaphase I, and 57% in metaphase II (Figure 6C-D). Addition of Bub1 kinase inhibitor did not significantly increase the number of oocytes with misaligned chromosomes in metaphase II but did show a small but statistically significant increase in metaphase I when compared to the controls. In contrast, inhibition of both haspin and Bub1 kinases resulted in 60% and 80% of oocytes with misaligned chromosomes in metaphase I and metaphase II, respectively. This percent of oocytes was similar to the percent of metaphase I and metaphase II oocytes with misaligned chromosomes with addition of the Aurora kinase inhibitor ZM447439 (Figure 6C-D). These results suggest that disruption of both recruitment pathways results in a loss of the pool of Aurora B/C kinase at the inner centromere required for proper chromosome alignment. Overall, our results support the model that haspin kinase has the most important role in error correction, but haspin and Bub1 recruitment pathways are compensatory for localization of Aurora B/C kinases for error correction of kinetochore-microtubule attachments in mouse oocytes.

## Discussion

In this study, we asked whether each pool of Aurora B kinase brought to the inner centromere and kinetochore by different recruitment pathways were important for the normal duration of anaphase onset and for error correction of kinetochore-microtubule attachments during meiosis and mitosis. By depleting or mutating the four different pathways in budding yeast, we first found that the pool of Ipl1^Aurora B^ brought by the Bub3/Bub1 pathway was the most important for delaying anaphase onset in meiosis (Figure 1C-E). For error correction of kinetochore-microtubule attachments, pools of Ipl1^Aurora B^ brought by both Bub3/Bub1 and the COMA complex had important individual roles in ensuring error correction during budding yeast meiosis (Figure 3D, I, J). In mitosis, the Bub3/Bub1 pathway and the COMA complex pathway are compensatory for ensuring bipolar kinetochore-microtubule attachments (Figure 4E). Finally, we asked if the Bub1/Bub3 and haspin kinase pathways had overlapping roles in recruiting AURKC for error correction of kinetochore-microtubule attachments during mammalian oogenesis. Inhibition of both haspin kinase and Bub1 kinase resulted in a strong increase in the number of oocytes with misaligned chromosomes, when compared to uninhibited controls and to oocytes where only one kinase was inhibited (Figure 6C, D). Our results suggest that the importance of each Aurora B/C recruitment pathway differs between mitosis and meiosis and among organisms.

### Differential Regulation of Budding Yeast Meiosis and Mitosis

One of the surprising aspects of our findings were the differences between mitosis and meiosis in the importance of each recruitment pathway for normal timing of anaphase onset and for error correction of kinetochore-microtubule attachments. Especially interesting is the differences between meiosis II and mitosis because both result in the separation of sister chromatids and were assumed to be regulated similarly. We show that loss of the pool of Ipl1^Aurora B^ brought by the Bub3/Bub1 pathway speeds up anaphase onset in meiosis II (Cairo et al., 2020; Yang et al., 2015)(Figure 1D). Loss of Bub3/Bub1 slows anaphase onset in mitosis (Yang et al., 2015)(Figure 1G). Furthermore, loss of either the Bub3/Bub1 recruitment pathway or the COMA complex recruitment pathway results in massive chromosome mis-segregation in meiosis II, whereas loss of both pathways are needed to cause massive chromosome mis-segregation in mitosis (Figure 3J, 4E) (Cairo et al., 2020; Yang et al., 2015).

To further understand how the pool of Ipl1^Aurora B^ brought by the Bub3/Bub1 pathway regulates the timing of anaphase onset, we investigated the role of PP1. Normally, Ipl1^Aurora B^ phosphorylates the kinetochore protein Knl1/Spc105 to prevent PP1 binding (Liu et al., 2010; Rosenberg et al., 2011). However, with depletion of Bub3 or Bub1, the levels of kinetochore-localized Ipl1^Aurora B^ were low, allowing premature PP1 binding (Cairo et al., 2020). Because previous studies showed that PP1 dephosphorylates the kinetochore-localized pool of Cdc20 for APC/C activation in the *C. elegans* embryo and in human cells, we considered that Cdc20 is also a PP1 substrate in budding yeast (Bancroft et al., 2020; Kim et al., 2017). S127 in Cdc20, identified in a phosphoproteomics study, is adjacent to a conserved sequence that contains a phosphorylation site important for regulating anaphase onset during *C. elegans* and human cell mitosis (Figure 2A) (Kim et al., 2017; Lanz et al., 2021). We mutated S127 to alanine and found a faster meiosis I and meiosis II onset (Figure 2B). Mutation of S88 and S89 to alanine, within a basic patch thought to bind PP1, significantly delayed anaphase I and anaphase II onset (Figure 2B). Interestingly, mitotic anaphase onset was not affected by the S127A, S88A,89A mutations (Figure 2E). These results suggest Cdc20’s dephosphorylation at the kinetochore in a PP1-dependent manner is sufficient to activate APC/C-Cdc20 in meiosis but not in mitosis.

In meiosis, our results support the model that anaphase onset timing is set by the balance of Ipl1^Aurora B^ and PP1 at the kinetochore. In this model, the pool of Ipl1^Aurora B^ brought by the Bub1/Bub3 recruitment pathway counteracts PP1 kinetochore binding. However, once bipolar attachments are established, Ipl1^Aurora B^ activity decreases and PP1 binds kinetochore protein Knl1/Spc105; PP1 then dephosphorylates kinetochore-localized Cdc20 for APC/C activation.

By studying the effect of different Cdc20 phosphomutants on setting the time for anaphase onset, we identified a remarkable difference between budding yeast mitotic and meiotic anaphase initiation regulation. We propose this difference between mitosis and meiosis anaphase initiation could be caused by a difference in the temporal regulation of Cdc20 kinetochore localization; in meiosis, Cdc20’s kinetochore localization or binding partners might make it more likely to be dephosphorylated in meiosis than in mitosis leading to APC/C-Cdc20 activation. Alternatively, or in addition, mitosis may have multiple layers of regulation, in that loss of a single recruitment pathway did not result in low levels of Ipl1^Aurora B^ at the kinetochore and phosphorylation status of Cdc20 at S127 is not important for anaphase onset timings (Figure 2E) (Cairo et al., 2020).

We previously showed less binding between APC/C and Cdc20 with loss of Bub3, which likely explains the anaphase onset delay in mitosis (Yang et al., 2015). The phosphorylation of other APC/C components, which are needed for Cdc20 binding, may be more important for the normal timing of anaphase onset in mitosis (Rudner and Murray, 2000). Whether the phosphorylation of these components occurs at the kinetochore is an intriguing future direction.

Also surprising was the difference between meiosis II and mitosis in the requirement for the Bub3/Bub1 and COMA complex recruitment pathways for normal chromosome segregation. In meiosis II, cells underwent massive chromosome mis-segregation with depletion of either recruitment pathway (Figure 3D, I-J). Most of the chromosomes stay attached to one SPB, leading to an uneven segregation of chromosomes due to a failure in correction of improper kinetochore-microtubule attachments (Cairo et al., 2020). In mitosis, the two pathways were compensatory; uneven chromosome segregation occurred only with depletion of both pathways (Figure 4E). These results suggest that both pools of Ipl1^Aurora B^ brought by Bub3/Bub1 and COMA complex are used in error correction of kinetochore-microtubule attachments. Furthermore, meiosis II may either require a higher level of kinetochore-localized Ipl1^Aurora B^ for error correction or pools of Ipl1^Aurora B^ at both the inner centromere and kinetochore.

### Error correction of kinetochore-microtubule attachments in mammalian oogenesis

Although addition of haspin or Bub1 kinase inhibitors resulted in decreased Aurora B kinetochore localization in mammalian cells, loss of either kinase activity did not have a major effect on chromosome segregation fidelity in a normal mitosis (Baron et al., 2016; Hadders et al., 2020). In contrast, loss of both haspin and Bub1 kinase activity results in an increase in lagging chromosomes, suggesting that the two pools are mostly redundant for their role in error correction (Hadders et al., 2020). However, the inhibition of Aurora B kinase has a much more severe defect than loss of both haspin and Bub1 kinase activity in mitotic cells, suggesting that another pool of Aurora B kinase is used for error correction of kinetochore-microtubule attachments. Indeed, several recent studies have identified a kinetochore-localized pool of Aurora B kinase, but the recruitment pathway needed for the localization is currently unknown (Broad et al., 2020; Hadders et al., 2020; Liang et al., 2020).

In mouse oocytes, Aurora C kinase is localized to the centromere and interchromatid axis (Avo Santos et al., 2011; Balboula et al., 2016; Tang et al., 2006; Uzbekova et al., 2008). Aurora B kinase, which is localized to the spindle, may also have a role in error correction (Balboula et al., 2017; Shuda et al., 2009). Inhibition of haspin kinase caused increased aneuploidy (Nguyen et al., 2014; Quartuccio et al., 2017). In contrast, loss of Bub1 kinase activity did not cause a severe defect in chromosome segregation (El Yakoubi et al., 2017; Ricke et al., 2012). We inhibited both haspin and Bub1 kinase to determine if the different pools of Aurora kinase C are both important for error correction. We monitored the alignment of chromosomes at the spindle equator in metaphase I and metaphase II to determine if there were misattachments (Figure 6A-B). Addition of haspin and Bub1 kinase inhibitors caused most oocytes to have misattached or misaligned chromosomes, similar to addition of Aurora B/C kinase inhibitor (Figure 6C-D).

Combined, these results suggest the model that although the pool of Aurora kinase C brought by haspin kinase is the most important, the pool brought by Bub1 kinase also has a role in the correction of improper kinetochore-microtubule attachments.

### Comparisons between yeast mouse oocytes

Our data suggests error correction of kinetochore-microtubule attachments in meiosis in yeast and mouse oocytes relies more heavily on individual recruitment pathways, however the combinational loss of multiple pathways gives more severe defects. In contrast, disruption of error correction in mitosis requires the loss of multiple recruitment pathways. This difference between meiosis and mitosis cannot be explained by the how chromosomes are segregated because both meiosis I and meiosis II have a greater reliance on the individual pools of Aurora kinases; and sister chromatid kinetochores are similarly positioned in both meiosis II and mitosis. We hypothesize that meiosis may require a higher level of kinetochore-localized Aurora B/C kinase for correction of inappropriate kinetochore-microtubule attachments. Alternatively, the distinct pools of Aurora kinase could each contribute differently to error correction between meiosis and mitosis.

Interestingly, the importance of the recruitment mechanisms differs between yeast and mouse. In yeast, the pool of Aurora B kinase brought by either the Bub3/Bub1 pathway or the COMA complex pathway is needed for proper chromosome segregation. In contrast, mouse oocytes rely most heavily on the pool of Aurora C kinase brought through the haspin kinase recruitment mechanism, with contribution of the pool brought through the Bub1 kinase mechanism. These results suggest that the importance of having multiple recruitment pathways has been maintained throughout evolution, but the reliance on each pathway has changed. Important future goals would be to further understand the mechanism of how each pool of Aurora B/C kinase interacts differently with kinetochore substrates in the presence and absence of tension.

## Materials and methods

### Budding yeast strains and manipulations

All *S. cerevisiae* strains used in this study are W303 derivatives and described in Table S1. All yeast strains contain the following genetic background: *ade2-1 his3-11,15 leu2-3,112 trp1-1 ura3-1* and *can1-100*. Standard PCR-based transformations were used to delete and tag genes (Janke et al., 2004). Genetic manipulations were checked by PCR, sequencing, and/or microscopy, if applicable. Strains with the anchor away background (*TOR1-1*, *fpr1Δ*, *RPL13-2XFKBP12*) were built as previously described but with *BUB3, IPL1, CTF19, SPC105* C-terminally tagged with FRB at the endogenous locus and *RPL13A* with *2XFKBP12*.

### Budding yeast growth conditions

For synchronized meiotic experiments, strains with *P_GAL1,10_-NDT80 Gal4-ER* were grown overnight in 2x synthetic complete medium (2xSC; 0.67% bacto–yeast nitrogen base without amino acids, 0.2% dropout mix with all amino acids, and 2% glucose) at 30⁰C, transferred with a 1:25 dilution to 2x synthetic complete acetate medium (2xSCA; 0.33% bacto-yeast nitrogen base without amino acids, 0.1% dropout mix with all amino acids, 1% potassium acetate) for 12-16h at 30⁰C, washed twice with water, and then incubated in 1% potassium acetate at 25⁰C for 11-12hrs to allow cells to reach prophase I arrest. After 11-12hrs in potassium acetate, β-estradiol (1μM; Sigma) was added. For anchor away in meiosis I and meiosis II, rapamycin (1μg/mL, Fisher BioReagents) was added 45min after β-estradiol addition. For anchor away in meiosis II, rapamycin was added 3hrs after β-estradiol addition. Time-lapse microscopy initiated ∼1h after β-estradiol addition.

For asynchronous meiotic experiments, cells were grown in 2xSC overnight at 30⁰C, transferred to 2xSCA (1:25) at 30⁰C for 12-16hrs, washed with water twice, and transferred to 1% potassium acetate at 25⁰C for 6-7hrs.

For asynchronous mitotic experiments, cells were grown overnight in 2xSC at 30⁰C and diluted (1:20). For anchor away experiments, rapamycin (1μg/mL, Fisher BioReagents) was added when diluting. Time-lapse microscopy was started 40-45min after dilution.

For synchronized mitotic experiments using strains in which the endogenous *CDC20* was under the methionine repressible *MET3* promoter (*P_MET_CDC20*), cells were grown in 2xSC medium lacking methionine overnight at 30⁰C. These cells were then diluted (1:20) in 2xSC medium lacking methionine and grown for 2hrs at 30⁰C. α-factor was added and cells were grown for 2 more hours at 30⁰C. After checking G1 arrest using microscopy, cells were washed 5-6 times with water and resuspended in 2xSC medium (containing methionine). Time-lapse microscopy was initiated 35-40min after washing out the α-factor.

All reagents used in this study are listed in Table S2.

### Microscope image acquisition and time-lapse microscopy in budding yeast

For time-lapse imaging, cells were concentrated and adhered to a coverslip coated with Concanavalin A (1mg/mL; Sigma; used 1xPBS as solvent) and inside a chamber. An agar pad (0.05g/mL) containing 1% potassium acetate (meiosis) or 2xSC (mitosis) was used to make a monolayer of cells and removed before imaging. Pre-conditioned 1% potassium acetate (meiosis) or 2xSC (mitosis) was added to the chamber before imaging. All meiosis movies were run for 12–13hrs and all mitosis movies were run for 6–8hrs.

*GFP-TUB1* and *SPC42-mCherry* expressing cells were imaged at room temperature using a Nikon Ti-E inverted-objective microscope equipped with a 60x oil objective (Plan Apochromat NA 1.4 oil), a Lambda 10–3 optical filter changer and Smartshutter, GFP and mCherry filters (Chroma Technology), and a charge-coupled device camera (CoolSNAP HQ2; Photometrics). During time-lapse imaging, z-stacks of five sections of 1.2µm each were acquired at 10-min intervals with exposure times of 60ms for bright field, 300-500ms for GFP, and 500-700 msec for mCherry with neutral-density filters transmitting 2–32% of light intensity.

*HTB2-mCherry* expressing cells were imaged at room temperature using a DeltaVision (pDV) microscope (Applied Precision) equipped with a CoolSNAP HQ2/HQ2-ICX285 camera using a 60× oil objective (U-Plan S-Apochromat-N, 1.4 NA). Images were acquired using SoftWoRx software (GE Healthcare). During time-lapse imaging, five z-steps (0.8µm) were acquired every 10 min. The exposure times used for bright-field and Htb2-mCherry were 0.3– 0.4msec and 0.02–0.03msec, respectively, with neutral-density filters transmitting 5–15% of light intensity.

*NDC80-yeGFP* expressing cells were imaged at room temperature using a DeltaVision Elite (eDV) microscope (Applied Precision) equipped with a PCO Edge5.5 sCMOS camera and Olympus 60X oil-immersion objective lens (Plan Apo N, 1.42). Images were acquired using SoftWoRx software (GE Healthcare). During time-lapse imaging, five z-steps (1µm) were acquired every 10 min. The exposure times used for bright-field, Spc42-mCherry, and Ndc80-yeGFP were 0.1msec, 0.5msec, and 0.25 msec, respectively, with neutral-density filters transmitting 32%, 10%, and 5% of light intensity, respectively.

### Image processing and quantification analysis in budding yeast

*GFP-TUB1* and *SPC42-mCherry* expressing cells that were imaged using a Nikon Ti-E inverted-objective microscope were analyzed using NIS-Elements software (Nikon). ImageJ was used to quantify images from mitotic and meiotic cells expressing *HTB2-mCherry*. To do so, we measured the area (μm^2^) of each DNA mass after the first and second meiotic divisions by adding all the area values in each z-stack (sum slices). To distinguish between even and uneven chromatin segregation, we measured the area of the DNA masses after mitosis and after the first and second meiotic division of our wildtype strains. For mitosis and meiosis I, the DNA mass area of the two DNA masses were subtracted from each other to get a mass size difference; similar calculations were done with the four DNA masses after meiosis II. The mass size difference values were averaged and the SD was used to define a range of ‘even’ chromatin segregation (average ± SD). For each individual cell, any mass size difference value outside the defined range from wildtype was classified as ‘uneven.’ The percent of cells with even and uneven chromatin segregation were plotted using Prism (GraphPad Software). Time-lapse images shown are the results of z-stacks combined into a single maximum intensity projection in Fiji (National Institutes of Health). Brightness and contrast were only adjusted on entire images.

*NDC80-yeGFP* fluorescence intensity at kinetochores (using colocalization with kinetochore protein *SPC42-mCherry*) in metaphase I was quantified with Fiji. To do so, a square was drawn around the *SPC42-mCherry* region and the sum intensity of the mCherry channel (A) and GFP channel (B) throughout the Z stacks was recorded. The background fluorescence for each channel, which was obtained by moving the circle for both mCherry (A_background_) and GFP (B_background_) to regions not containing *SPC42-mCherry* or *NDC80-yeGFP*, was subtracted. To determine the fluorescence intensity of *NDC80-yeGFP* at the kinetochore, we calculated the ratio between *NDC80-yeGFP* and *SPC42-mCherry* as follows: (B-B_background_)/(A-A_background_). All ratio values were plotted using Prism (GraphPad Software). Images shown are the results of z-stacks combined into a single maximum intensity projection in Fiji (National Institutes of Health). Brightness and contrast were only adjusted on entire images.

### Oocyte collection and culture

We used wildtype CF1 mice from Envigo Laboratories for mouse experiments. Mice were housed in 12-12hr light-dark cycle, with constant temperature, food, and water. All oocyte experiments were conducted using healthy female mice ranging from 22-24 grams (6-8 weeks in age). Mice were not involved in other experimental procedures. Prophase I arrested oocytes were collected from the ovaries of female wildtype mice using minimal essential medium (MEM) (MEM with Earle’s Salts, 0.1g/L pyruvate, 25 mM HEPES pH 7.3, 3 mg/mL poyvinylpyrrolidone). To mature oocytes, we used Chatot, Ziomek, and Bavister (CZB) medium (81.6mM NaCl, 4.8mM KCl, 1.2mM KH_2_PO_4_, 1.2mM MgSO_4_ 7H_2_O, 0.27mM pyruvic acid, 1.7mM CaCl_2_ 2H_2_O, 30.8mM DL-Lactic acid, 7mM Taurine, 0.1mM EDTA, 25mM NaHCO_3_, gentamicin 1:1,000, 1:1,000 1% phenol red, 300mg/mL BSA) with fresh glutamine (1mM, Sigma) and incubated the oocytes at 37⁰C with 5% CO_2_^4^. While collecting oocytes, milrinone (Sigma Aldrich, 2.5mM) was added to MEM to prevent meiotic resumption, which was subsequently washed with CZB to allow oocytes to mature. To better visualize kinetochores and Aurora kinase C in mouse oocytes, we removed the zona pellucida by washing the oocytes in 4-6 drops of acidic Tyrode’s solution (Millipore Sigma) and rinsed them in 4-6 drops of CZB media with the corresponding drug(s) and/or solvents prior to fixation. All reagents used in this study are listed in Table S2.

### Kinase Inhibitor Addition

All inhibitors used for the mouse oocytes are listed in Table S3. 5-Iodotubercidin (5-Itu; Cayman Chemicals; 0.5μM) was used to inhibit haspin kinase with ethanol (Fisher) as the control (solvent for 5-Itu). To disrupt Bub1 H2A phosphorylation, we added BAY-1816032 (8μM, MedChem Express); we used DMSO (Sigma) as control (solvent for BAY-1816032). We used ZM447439 (Tocris Bioscience, 5μM) as an Aurora B/C inhibitor and DMSO (Sigma) as a control (solvent for ZM447439). For all mouse meiosis I experiments, BAY-1816032 and ZM447439 were added at 0h when transferring oocytes to CZB, while 5-Itu was added after 5hrs from CZB oocyte transfer. For all mouse meiosis II experiments, all drugs were added at 11hrs. Oocytes were fixed 8hrs after CZB oocyte transfer for meiosis I experiments and after 15hrs for meiosis II experiments. We treated all meiosis I mouse oocytes with acidic Tyrode’s solution (Millipore Sigma) to remove the zone pellucida when imaging for Aurora kinase C, kinetochores, and DNA.

### Mouse oocyte immunofluorescence

Oocytes were fixed in phosphate-buffered saline (PBS) (2.7mM KCl, 1.8mM KH_2_PO_4_, 137mM NaCl, 10mM Na_2_HPO_4_ 7H_2_O) containing 2% PFA for 20min at room temperature for detection of microtubules, kinetochores, and Aurora kinase C. For detection of H2AT120ph and kinetochores, oocytes were fixed in 3.7% PFA containing 0.1% Triton X-100 for 1h at room temperature. Fixed oocytes were transferred to blocking solution (PBS, 0.3% BSA wt/vol, 0.01% Tween-20 wt/vol) at 4°C overnight. Prior to immunofluorescence staining, oocytes were transferred to permeabilization solution (PBS, 0.3% wt/vol, 0.1% vol/vol Triton X-100) for 20min and later washed with blocking solution 3 times. Next, oocytes were incubated in primary antibody diluted in blocking solution for 1h at room temperature inside a humidified chamber.

After washing with blocking solution 3 times for 10min, oocytes were incubated with the corresponding secondary antibody for 1h at room temperature inside the humidified chamber. After washing with blocking solution 3 times for 10 min, oocytes were mounted in 5μL Vectashield (Vector Laboratories) containing DAPI (Thermo Fisher Scientific, 0.15mg/mL) to visualize DNA.

To visualize microtubules, we used rabbit monoclonal anti-α-tubulin conjugated with Alexa Fluor 488 (1:30, Cell Signaling). To detect kinetochores, we used human polyclonal anti-CREST/ACA (1:15; Antibodies Incorporated) antibody and goat anti-human Alexa-Fluor 633 (Life Technologies, 1:200). To visualize Aurora kinase C, we used rabbit anti-mAurC (1:500, FORTIS) and Alexa-Fluor 568 donkey anti-rabbit IgG (1:200, Invitrogen). For H2AT120ph, we used Histone H2AT120ph pAb (1:500, Active Motif) and goat anti-rabbit Alexa-Fluor 488 (Life Technologies, 1:200). All antibodies used for mouse work are listed in Table S4.

### Imaging mouse oocytes

All mouse oocytes were imaged at room temperature using a DeltaVision pDV microscope (Applied Precision) equipped with a CoolSNAP HQ2/HQ2-ICX285 camera using a 100x oil objective (U-Plan S-Apochromat-N, 1.4 NA). Images were acquired using SoftWoRx software (GE Heathcare). To visualize microtubules (0.025msec, 2% transmittance) and DNA (0.025 msec, 2% transmittance), z-stacks were taken 0.4µm apart. To visualize Aurora kinase C (0.25msec, 5% transmittance), kinetochores (0.8msec, 10% transmittance), and DNA (0.2msec, 2% transmittance), z-stacks were taken 0.2µm apart. To visualize H2AT120ph (0.08msec, 2% transmittance), kinetochores (0.25msec, 50% transmittance), and DNA (0.025 msec, 2% transmittance), z-stacks were taken 0.2µm apart.

### Image processing and quantification analysis in mouse oocytes

We processed all images using ImageJ. Z-stacks were combined into a single maximum intensity projection to obtain the final images presented in this article. Brightness and contrast were only adjusted on entire images. Aurora kinase C and H2AT120ph fluorescence intensity at kinetochores was quantified with ImageJ. To do so, a circle was drawn around the kinetochore region and the intensity of the kinetochore channel (A) and Aurora kinase C or H2AT120ph channel (B) was recorded. The background fluorescence for each channel, which was obtained by moving the circle for both channels to regions not containing kinetochores (A_background_ and B_background_), was subtracted. To determine the fluorescence intensity of Aurora kinase C and H2AT120ph at kinetochores, we calculated the following ratio in each case: (B-B_background_)/(A-A_background_). All ratio values were plotted using Prism (GraphPad Software).

### Statistical analysis

Statistical analysis was performed in Graphpad Prism. For yeast anaphase onset timing and all yeast and mouse fluorescence intensity measurements, an unpaired, nonparametric Mann-Whitney test with computation of two-tailed exact P values was used. The two-sided Fisher’s exact test was used to analyze both the yeast DNA masses (Htb2-mCherry strains) and the mouse chromosome attachment/alignment percentages.

## Acknowledgements

We thank the Lacefield lab for insightful comments on the manuscript. We thank Trisha Davis and Andreas Hochwagen for strains. We thank the Light Microscopy and Imaging Center at Indiana University, especially Jim Powers for assistance. This research was supported by NIH grant GM105755 to SL and NIH grants R35GM136340 and R01 GM112801 to KS.

## Declaration of Interests

The authors do not have any competing interests to declare.

## Supplementary Figure Legends

**Figure S1.**
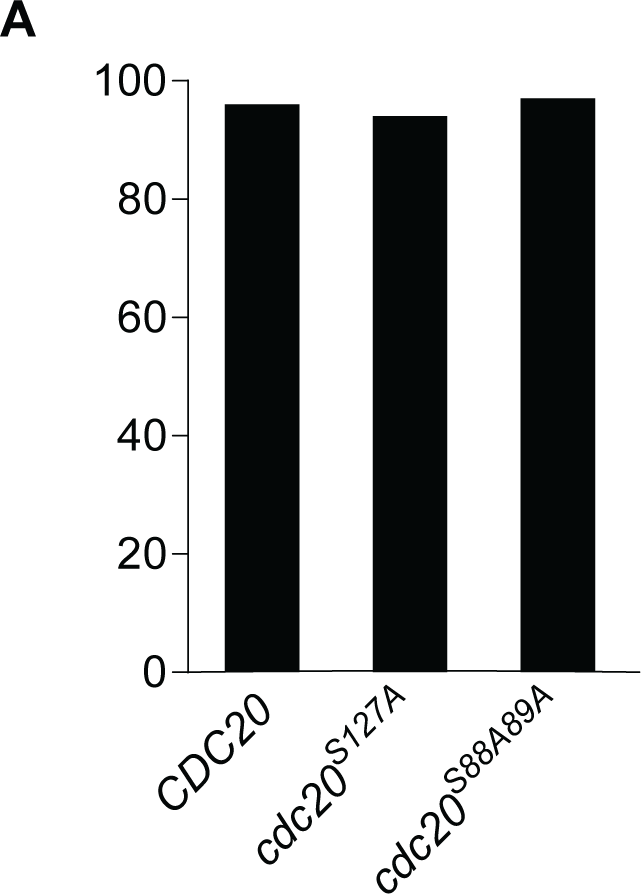
Spore viability Cdc20 phosphomutants. Related to Figure 2. (A) Percent sporulation of the indicated genotypes. All strains have the endogenous *CDC20* under the mitosis-specific CLB2 promoter and express an integrated *CDC20*, *cdc20^S127A^* or *cdc20^S88A,S89A^* under the *CDC20* promoter. Data were obtained by analyzing the growth of ≥240 spores for each genotype.

**Figure S2.**
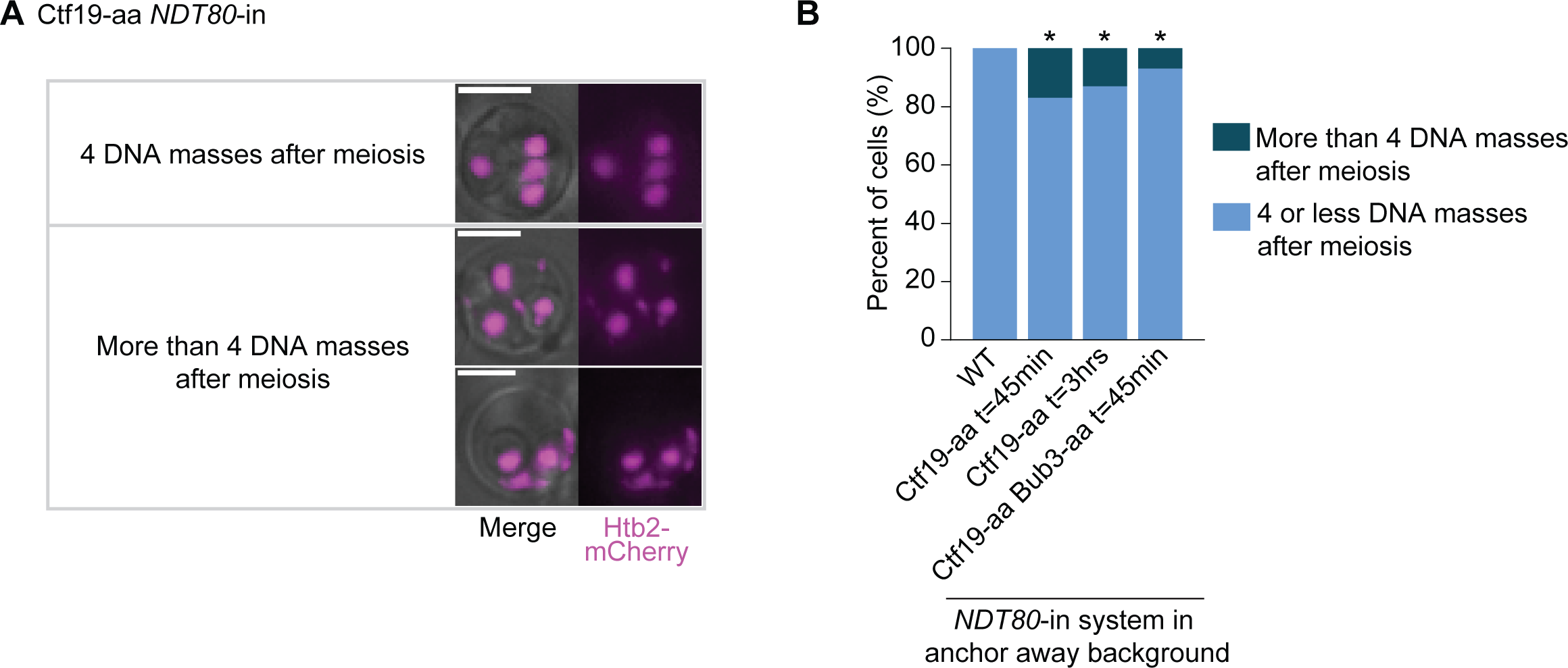
Ctf19 nuclear depletion prior to prophase I results in additional DNA masses after meiosis. Related to Figure 3. (A) Representative images of Ctf19-aa *NDT80*-in cells upon rapamycin addition 3 hours after β-estradiol release displaying 4 or more DNA masses after meiosis. Cells express Htb2-mCherry to visualize chromatin. Scale bars, 5μm. (B) Percent of cells that display less than 4, 4, or more than 4 DNA masses after meiosis, containing the *NDT80*-in (*P_GAL1,10_-NDT80*, *Gal4-ER*) system and anchor away background (*tor1-1*, *fpr1Δ*, *RPL13-2XFKBP12*). In all strains, β-estradiol was added 12 hours after cells were resuspended in sporulation media. As indicated, rapamycin was added to strains containing Bub3-aa and Ctf19-aa 45 minutes or 3hrs after β-estradiol addition to deplete Bub3 and Ctf19 from the nucleus, respectively. * indicates a statistically significant difference compared to wildtype anchor away cells (at least 100 cells from two or more independent experiments per genotype were counted; P < 0.05, two-tailed Fisher’s exact test). aa, anchor away.

**Figure S3.**
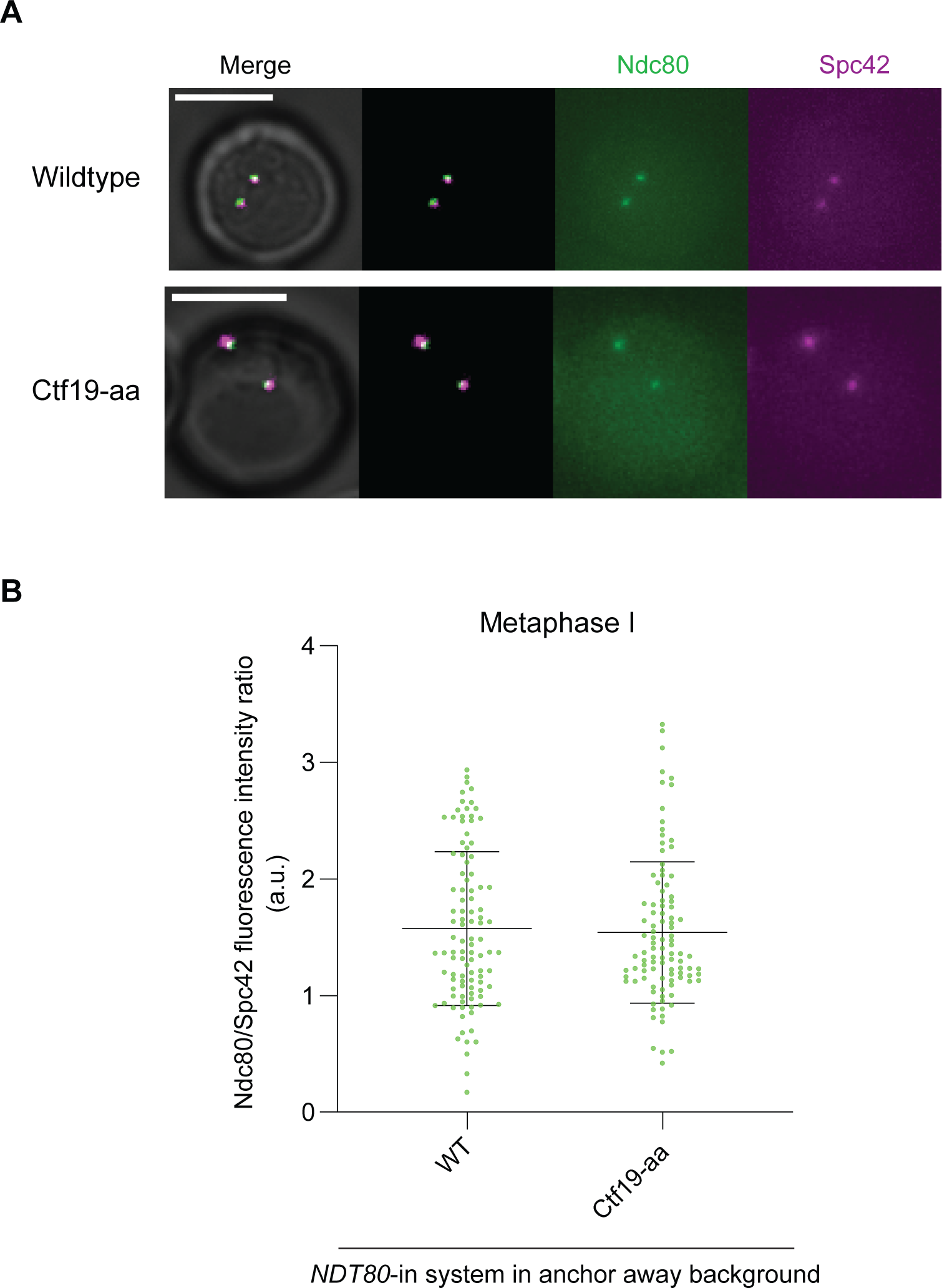
Ctf19 nuclear depletion 45 minutes after pachytene release does not affect kinetochore integrity in metaphase I. Related to Figure 3. (A) Representative images of Ctf19-aa *NDT80*-in cells upon rapamycin addition 45 minutes after β-estradiol release. Cells express *NDC80-yeGFP* and *SPC42-mCherry* to visualize Ndc80 and Spc42. Scale bars, 5μm. (B) Graph indicating Ndc80 fluorescence intensity relative to kinetochores (Spc42). Each point plotted represents a single kinetochore. Strains contain the *NDT80*-in system (*P_GAL1,10_-NDT80*, *Gal4-ER*) and the anchor away genetic background (*TOR1-1*, *fpr1Δ*, *RPL13-2XFKBP12*). In all strains, β-estradiol was added 12 hours after cells were resuspended in sporulation media. Rapamycin was added 45 minutes after β-estradiol addition to deplete Ctf19 from the nucleus. 100 cells from per genotype were counted; P < 0.05, Mann-Whitney test; error bars show SD). aa, anchor away.

**Figure S4.**
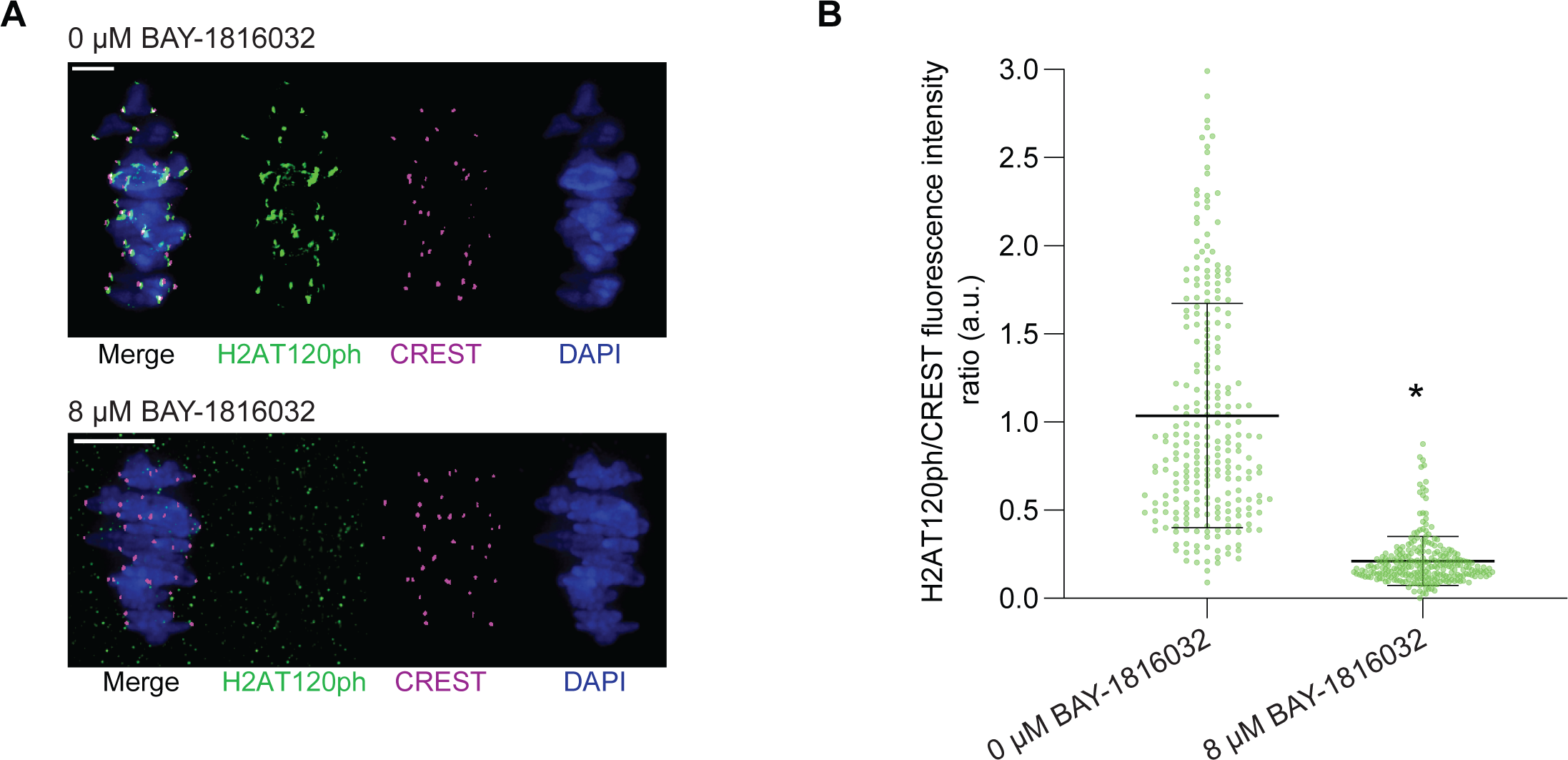
8μM BAY-1816032 abolishes H2AT120ph staining at kinetochores. Related to Figure 5 and 6. (A) Representative images of metaphase I oocytes treated with either 0μM (DMSO) or 8μM of BAY-1816032. Oocytes were matured *in vitro* for 8 hours and immunostained for H2AT120ph (green), kinetochores (CREST, magenta), and DNA (DAPI, blue). DMSO or BAY-1816032 were added when oocytes were transferred to the maturation media (CZB) during prophase I. Scale bars, 10μm. (B) Graph indicating H2AT120ph fluorescence intensity relative to kinetochores (CREST) staining (H2AT120ph/CREST). Each point plotted represents a single kinetochore. 25 kinetochores were scored in each oocyte. 10 metaphase I oocytes were used for each concentration. P<0.05, Mann-Whitney test; dot plot shows SD.

## Notes

### Competing Interest Statement

The authors have declared no competing interest.

## References

Abad, M. A., Gupta, T., Hadders, M. A., Meppelink, A., Wopken, J. P., Blackburn, E., Zou, J., Gireesh, A., Buzuk, L., Kelly, D. A. et al. (2022). Mechanistic basis for Sgo1-mediated centromere localization and function of the CPC. J Cell Biol 221.

Avo Santos, M., van de Werken, C., de Vries, M., Jahr, H., Vromans, M. J., Laven, J. S., Fauser, B. C., Kops, G. J., Lens, S. M. and Baart, E. B. (2011). A role for Aurora C in the chromosomal passenger complex during human preimplantation embryo development. Hum Reprod 26, 1868–81.

Balboula, A. Z., Blengini, C. S., Gentilello, A. S., Takahashi, M. and Schindler, K. (2017). Maternal RNA regulates Aurora C kinase during mouse oocyte maturation in a translation-independent fashion. Biol Reprod 96, 1197–1209.

Balboula, A. Z., Nguyen, A. L., Gentilello, A. S., Quartuccio, S. M., Drutovic, D., Solc, P. and Schindler, K. (2016). Haspin kinase regulates microtubule-organizing center clustering and stability through Aurora kinase C in mouse oocytes. J Cell Sci 129, 3648–3660.

Bancroft, J., Holder, J., Geraghty, Z., Alfonso-Perez, T., Murphy, D., Barr, F. A. and Gruneberg, U. (2020). PP1 promotes cyclin B destruction and the metaphase-anaphase transition by dephosphorylating CDC20. Mol Biol Cell 31, 2315–2330.

Baron, A. P., von Schubert, C., Cubizolles, F., Siemeister, G., Hitchcock, M., Mengel, A., Schroder, J., Fernandez-Montalvan, A., von Nussbaum, F., Mumberg, D. et al. (2016). Probing the catalytic functions of Bub1 kinase using the small molecule inhibitors BAY-320 and BAY-524. Elife 5.

Benjamin, K. R., Zhang, C., Shokat, K. M. and Herskowitz, I. (2003). Control of landmark events in meiosis by the CDK Cdc28 and the meiosis-specific kinase Ime2. Genes Dev 17, 1524–39.

Borek, W. E., Vincenten, N., Duro, E., Makrantoni, V., Spanos, C., Sarangapani, K. K., de Lima Alves, F., Kelly, D. A., Asbury, C. L., Rappsilber, J. et al. (2021). The Proteomic Landscape of Centromeric Chromatin Reveals an Essential Role for the Ctf19(CCAN) Complex in Meiotic Kinetochore Assembly. Curr Biol 31, 283–296 e7.

Broad, A. J., DeLuca, K. F. and DeLuca, J. G. (2020). Aurora B kinase is recruited to multiple discrete kinetochore and centromere regions in human cells. J Cell Biol 219.

Cairo, G. and Lacefield, S. (2020). Establishing correct kinetochore-microtubule attachments in mitosis and meiosis. Essays Biochem 64, 277–287.

Cairo, G., MacKenzie, A., Tsuchiya, D. and Lacefield, S. (2022). Use of Time-Lapse Microscopy and Stage-Specific Nuclear Depletion of Proteins to Study Meiosis in S. Cerevisiae. J Vis Exp.

Cairo, G., MacKenzie, A. M. and Lacefield, S. (2020). Differential requirement for Bub1 and Bub3 in regulation of meiotic versus mitotic chromosome segregation. J Cell Biol 219.

Campbell, C. S. and Desai, A. (2013). Tension sensing by Aurora B kinase is independent of survivin-based centromere localization. Nature 497, 118–21.

Carlile, T. M. and Amon, A. (2008). Meiosis I is established through division-specific translational control of a cyclin. Cell 133, 280–91.

Cho, U. S. and Harrison, S. C. (2011). Ndc10 is a platform for inner kinetochore assembly in budding yeast. Nat Struct Mol Biol 19, 48–55.

Dahmann, C. and Futcher, B. (1995). Specialization of B-type cyclins for mitosis or meiosis in S. cerevisiae. Genetics 140, 957–63.

Dai, J., Sultan, S., Taylor, S. S. and Higgins, J. M. (2005). The kinase haspin is required for mitotic histone H3 Thr 3 phosphorylation and normal metaphase chromosome alignment. Genes Dev 19, 472–88.

Edgerton, H., Johansson, M., Keifenheim, D., Mukherjee, S., Chacon, J. M., Bachant, J., Gardner, M. K. and Clarke, D. J. (2016). A noncatalytic function of the topoisomerase II CTD in Aurora B recruitment to inner centromeres during mitosis. J Cell Biol 213, 651–64.

El Yakoubi, W., Buffin, E., Cladiere, D., Gryaznova, Y., Berenguer, I., Touati, S. A., Gomez, R., Suja, J. A., van Deursen, J. M. and Wassmann, K. (2017). Mps1 kinase-dependent Sgo2 centromere localisation mediates cohesin protection in mouse oocyte meiosis I. Nat Commun 8, 694.

Fischbock-Halwachs, J., Singh, S., Potocnjak, M., Hagemann, G., Solis-Mezarino, V., Woike, S., Ghodgaonkar-Steger, M., Weissmann, F., Gallego, L. D., Rojas, J. et al. (2019). The COMA complex interacts with Cse4 and positions Sli15/Ipl1 at the budding yeast inner kinetochore. Elife 8.

Funk, L. C., Zasadil, L. M. and Weaver, B. A. (2016). Living in CIN: Mitotic Infidelity and Its Consequences for Tumor Promotion and Suppression. Dev Cell 39, 638–652.

Galli, M., Diani, L., Quadri, R., Nespoli, A., Galati, E., Panigada, D., Plevani, P. and Muzi-Falconi, M. (2020). Haspin Modulates the G2/M Transition Delay in Response to Polarization Failures in Budding Yeast. Front Cell Dev Biol 8, 625717.

Garcia-Rodriguez, L. J., Kasciukovic, T., Denninger, V. and Tanaka, T. U. (2019). Aurora B-INCENP Localization at Centromeres/Inner Kinetochores Is Required for Chromosome Bi-orientation in Budding Yeast. Curr Biol 29, 1536–1544 e4.

Gassmann, R., Carvalho, A., Henzing, A. J., Ruchaud, S., Hudson, D. F., Honda, R., Nigg, E. A., Gerloff, D. L. and Earnshaw, W. C. (2004). Borealin: a novel chromosomal passenger required for stability of the bipolar mitotic spindle. J Cell Biol 166, 179–91.

Hadders, M. A., Hindriksen, S., Truong, M. A., Mhaskar, A. N., Wopken, J. P., Vromans, M. J. M. and Lens, S. M. A. (2020). Untangling the contribution of Haspin and Bub1 to Aurora B function during mitosis. J Cell Biol 219.

Haruki, H., Nishikawa, J. and Laemmli, U. K. (2008). The anchor-away technique: rapid, conditional establishment of yeast mutant phenotypes. Mol Cell 31, 925–32.

Hendrickx, A., Beullens, M., Ceulemans, H., Den Abt, T., Van Eynde, A., Nicolaescu, E., Lesage, B. and Bollen, M. (2009). Docking motif-guided mapping of the interactome of protein phosphatase-1. Chem Biol 16, 365–71.

Heroes, E., Lesage, B., Gornemann, J., Beullens, M., Van Meervelt, L. and Bollen, M. (2013). The PP1 binding code: a molecular-lego strategy that governs specificity. FEBS J 280, 584–95.

Higgins, J. M. (2003). Structure, function and evolution of haspin and haspin-related proteins, a distinctive group of eukaryotic protein kinases. Cell Mol Life Sci 60, 446–62.

Hyland, K. M., Kingsbury, J., Koshland, D. and Hieter, P. (1999). Ctf19p: A novel kinetochore protein in Saccharomyces cerevisiae and a potential link between the kinetochore and mitotic spindle. J Cell Biol 145, 15–28.

Janke, C., Magiera, M. M., Rathfelder, N., Taxis, C., Reber, S., Maekawa, H., Moreno-Borchart, A., Doenges, G., Schwob, E., Schiebel, E. et al. (2004). A versatile toolbox for PCR-based tagging of yeast genes: new fluorescent proteins, more markers and promoter substitution cassettes. Yeast 21, 947–62.

Jeyaprakash, A. A., Basquin, C., Jayachandran, U. and Conti, E. (2011). Structural basis for the recognition of phosphorylated histone h3 by the survivin subunit of the chromosomal passenger complex. Structure 19, 1625–34.

Kawashima, S. A., Yamagishi, Y., Honda, T., Ishiguro, K. and Watanabe, Y. (2010). Phosphorylation of H2A by Bub1 prevents chromosomal instability through localizing shugoshin. Science 327, 172–7.

Kelly, A. E., Ghenoiu, C., Xue, J. Z., Zierhut, C., Kimura, H. and Funabiki, H. (2010). Survivin reads phosphorylated histone H3 threonine 3 to activate the mitotic kinase Aurora B. Science 330, 235–9.

Kim, T., Lara-Gonzalez, P., Prevo, B., Meitinger, F., Cheerambathur, D. K., Oegema, K. and Desai, A. (2017). Kinetochores accelerate or delay APC/C activation by directing Cdc20 to opposing fates. Genes Dev 31, 1089–1094.

Kitajima, T. S., Hauf, S., Ohsugi, M., Yamamoto, T. and Watanabe, Y. (2005). Human Bub1 defines the persistent cohesion site along the mitotic chromosome by affecting Shugoshin localization. Curr Biol 15, 353–9.

Kitajima, T. S., Sakuno, T., Ishiguro, K., Iemura, S., Natsume, T., Kawashima, S. A. and Watanabe, Y. (2006). Shugoshin collaborates with protein phosphatase 2A to protect cohesin. Nature 441, 46–52.

Krenn, V. and Musacchio, A. (2015). The Aurora B Kinase in Chromosome Bi-Orientation and Spindle Checkpoint Signaling. Front Oncol 5, 225.

Lanz, M. C., Yugandhar, K., Gupta, S., Sanford, E. J., Faca, V. M., Vega, S., Joiner, A. M. N., Fromme, J. C., Yu, H. and Smolka, M. B. (2021). In-depth and 3-dimensional exploration of the budding yeast phosphoproteome. EMBO Rep 22, e51121.

Liang, C., Zhang, Z., Chen, Q., Yan, H., Zhang, M., Zhou, L., Xu, J., Lu, W. and Wang, F. (2020). Centromere-localized Aurora B kinase is required for the fidelity of chromosome segregation. J Cell Biol 219.

Liu, D., Vleugel, M., Backer, C. B., Hori, T., Fukagawa, T., Cheeseman, I. M. and Lampson, M. A. (2010). Regulated targeting of protein phosphatase 1 to the outer kinetochore by KNL1 opposes Aurora B kinase. J Cell Biol 188, 809–20.

Liu, H., Jia, L. and Yu, H. (2013a). Phospho-H2A and cohesin specify distinct tension-regulated Sgo1 pools at kinetochores and inner centromeres. Curr Biol 23, 1927–33.

Liu, H., Qu, Q., Warrington, R., Rice, A., Cheng, N. and Yu, H. (2015). Mitotic Transcription Installs Sgo1 at Centromeres to Coordinate Chromosome Segregation. Mol Cell 59, 426–36.

Liu, H., Rankin, S. and Yu, H. (2013b). Phosphorylation-enabled binding of SGO1-PP2A to cohesin protects sororin and centromeric cohesion during mitosis. Nat Cell Biol 15, 40–9.

MacKenzie, A. M. and Lacefield, S. (2020). CDK Regulation of Meiosis: Lessons from S. cerevisiae and S. pombe. Genes (Basel) 11.

Meyer, R. E., Kim, S., Obeso, D., Straight, P. D., Winey, M. and Dawson, D. S. (2013).

Mps1 and Ipl1/Aurora B act sequentially to correctly orient chromosomes on the meiotic spindle of budding yeast. *Science* 339, 1071-4.

Nagaoka, S. I., Hassold, T. J. and Hunt, P. A. (2012). Human aneuploidy: mechanisms and new insights into an age-old problem. Nat Rev Genet 13, 493–504.

Nespoli, A., Vercillo, R., di Nola, L., Diani, L., Giannattasio, M., Plevani, P. and Muzi-Falconi, M. (2006). Alk1 and Alk2 are two new cell cycle-regulated haspin-like proteins in budding yeast. Cell Cycle 5, 1464–71.

Nguyen, A. L., Gentilello, A. S., Balboula, A. Z., Shrivastava, V., Ohring, J. and Schindler,

K. (2014). Phosphorylation of threonine 3 on histone H3 by haspin kinase is required for meiosis I in mouse oocytes. J Cell Sci 127, 5066–78.

Ortiz, J., Stemmann, O., Rank, S. and Lechner, J. (1999). A putative protein complex consisting of Ctf19, Mcm21, and Okp1 represents a missing link in the budding yeast kinetochore. Genes Dev 13, 1140–55.

Panigada, D., Grianti, P., Nespoli, A., Rotondo, G., Castro, D. G., Quadri, R., Piatti, S., Plevani, P. and Muzi-Falconi, M. (2013). Yeast haspin kinase regulates polarity cues necessary for mitotic spindle positioning and is required to tolerate mitotic arrest. Dev Cell 26, 483–95.

Quartuccio, S. M., Dipali, S. S. and Schindler, K. (2017). Haspin inhibition reveals functional differences of interchromatid axis-localized AURKB and AURKC. Mol Biol Cell 28, 2233–2240.

Ricke, R. M., Jeganathan, K. B., Malureanu, L., Harrison, A. M. and van Deursen, J. M. (2012). Bub1 kinase activity drives error correction and mitotic checkpoint control but not tumor suppression. J Cell Biol 199, 931–49.

Rosenberg, J. S., Cross, F. R. and Funabiki, H. (2011). KNL1/Spc105 recruits PP1 to silence the spindle assembly checkpoint. Curr Biol 21, 942–7.

Rudner, A. D. and Murray, A. W. (2000). Phosphorylation by Cdc28 activates the Cdc20-dependent activity of the anaphase-promoting complex. J Cell Biol 149, 1377–90.

Sampath, S. C., Ohi, R., Leismann, O., Salic, A., Pozniakovski, A. and Funabiki, H. (2004). The chromosomal passenger complex is required for chromatin-induced microtubule stabilization and spindle assembly. Cell 118, 187–202.

Saurin, A. T. (2018). Kinase and Phosphatase Cross-Talk at the Kinetochore. Front Cell Dev Biol 6, 62.

Shuda, K., Schindler, K., Ma, J., Schultz, R. M. and Donovan, P. J. (2009). Aurora kinase B modulates chromosome alignment in mouse oocytes. Mol Reprod Dev 76, 1094–105.

Siemeister, G., Mengel, A., Fernandez-Montalvan, A. E., Bone, W., Schroder, J., Zitzmann-Kolbe, S., Briem, H., Prechtl, S., Holton, S. J., Monning, U. et al. (2019). Inhibition of BUB1 Kinase by BAY 1816032 Sensitizes Tumor Cells toward Taxanes, ATR, and PARP Inhibitors In Vitro and In Vivo. Clin Cancer Res 25, 1404–1414.

Smith, R. J., Cordeiro, M. H., Davey, N. E., Vallardi, G., Ciliberto, A., Gross, F. and Saurin, A. T. (2019). PP1 and PP2A Use Opposite Phospho-dependencies to Control Distinct Processes at the Kinetochore. Cell Rep 28, 2206–2219 e8.

Tanaka, T. U., Rachidi, N., Janke, C., Pereira, G., Galova, M., Schiebel, E., Stark, M. J. and Nasmyth, K. (2002). Evidence that the Ipl1-Sli15 (Aurora kinase-INCENP) complex promotes chromosome bi-orientation by altering kinetochore-spindle pole connections. Cell 108, 317–29.

Tang, C. J., Lin, C. Y. and Tang, T. K. (2006). Dynamic localization and functional implications of Aurora-C kinase during male mouse meiosis. Dev Biol 290, 398–410.

Tsuchiya, D. and Lacefield, S. (2013). Cdk1 modulation ensures the coordination of cell-cycle events during the switch from meiotic prophase to mitosis. Curr Biol 23, 1505–13.

Tsukahara, T., Tanno, Y. and Watanabe, Y. (2010). Phosphorylation of the CPC by Cdk1 promotes chromosome bi-orientation. Nature 467, 719–23.

Uhlmann, F., Wernic, D., Poupart, M. A., Koonin, E. V. and Nasmyth, K. (2000). Cleavage of cohesin by the CD clan protease separin triggers anaphase in yeast. Cell 103, 375–86.

Uzbekova, S., Arlot-Bonnemains, Y., Dupont, J., Dalbies-Tran, R., Papillier, P., Pennetier, S., Thelie, A., Perreau, C., Mermillod, P., Prigent, C. et al. (2008). Spatio-temporal expression patterns of aurora kinases a, B, and C and cytoplasmic polyadenylation-element-binding protein in bovine oocytes during meiotic maturation. Biol Reprod 78, 218–33.

Wang, F., Dai, J., Daum, J. R., Niedzialkowska, E., Banerjee, B., Stukenberg, P. T., Gorbsky, G. J. and Higgins, J. M. (2010). Histone H3 Thr-3 phosphorylation by Haspin positions Aurora B at centromeres in mitosis. Science 330, 231–5.

Widlund, P. O., Lyssand, J. S., Anderson, S., Niessen, S., Yates, J. R., 3rd and Davis, T. N. (2006). Phosphorylation of the chromosomal passenger protein Bir1 is required for localization of Ndc10 to the spindle during anaphase and full spindle elongation. Mol Biol Cell 17, 1065–74.

Xu, L., Ajimura, M., Padmore, R., Klein, C. and Kleckner, N. (1995). NDT80, a meiosis-specific gene required for exit from pachytene in Saccharomyces cerevisiae. Mol Cell Biol 15, 6572–81.

Xu, Z., Cetin, B., Anger, M., Cho, U. S., Helmhart, W., Nasmyth, K. and Xu, W. (2009). Structure and function of the PP2A-shugoshin interaction. Mol Cell 35, 426–41.

Yamagishi, Y., Honda, T., Tanno, Y. and Watanabe, Y. (2010). Two histone marks establish the inner centromere and chromosome bi-orientation. Science 330, 239–43.

Yang, Y., Tsuchiya, D. and Lacefield, S. (2015). Bub3 promotes Cdc20-dependent activation of the APC/C in S. cerevisiae. J Cell Biol 209, 519–27.

Yoon, H. J. and Carbon, J. (1999). Participation of Bir1p, a member of the inhibitor of apoptosis family, in yeast chromosome segregation events. Proc Natl Acad Sci U S A 96, 13208–13.

