## Supplemental Tables for "Distinct Aurora B pools at the inner centromere and kinetochore have different contributions to meiotic and mitotic chromosome segregation"

**Table S1. Budding yeast strains used in this study**

| Strain number: | Strain genotype: |
|---|---|
| LY3565 | *MATa/α, $P_{GAL1,10}$-NDT80:TRP1/ $P_{GAL1,10}$-NDT80:TRP1, Gal4-ER:URA3/Gal4-ER:URA3, Spc42-mCherry:kanMX6/+, $P_{TUB1}$GFP-TUB1:LEU2/+, ZIP1-GFP/+* |
| LY7943 | *MATa/α, $P_{GAL1,10}$-NDT80:TRP1/ $P_{GAL1,10}$-NDT80:TRP1, Gal4-ER:URA3/Gal4-ER:URA3, Spc42-mCherry:kanMX6/+, $P_{TUB1}$GFP-TUB1:LEU2/+, 12myc-cdc20/12myc-cdc20* |
| LY5417 | *MATa*, *Spc42-mCherry: hphNT1/+, $P_{TUB1}$GFP-TUB1:URA3/+*, *bar1::TRP1* |
| LY8488 | *MATa*, *Spc42-mCherry: hphNT1/+, $P_{TUB1}$GFP-TUB1:URA3/+, bar1::TRP1*, *12myc-cdc20/12myc-cdc20* |
| LY3022 | *MATa/α, Spc42-mCherry: hphNT1/+, $P_{TUB1}$GFP-TUB1:URA3/ $P_{TUB1}$GFP-TUB1:URA3*, *ZIP1-GFP/+* |
| LY3617 | *MATa/α, tor1-1::HIS3/tor1-1, fpr1::loxP-LEU2-loxP/fpr1::natMX4, RPL13A-2XFKBP12::loxP/RPL13A-2XFKBP12::TRP1, Spc42-mCherry:hphNT1/+, $P_{TUB1}$GFP-TUB1:URA3/+, ZIP1-GFP/+* |
| LY8258 | *MATa/α, spc105-RVAF/spc105-RVAF, Spc42-mCherry: hphNT1/+, $P_{TUB1}$GFP-TUB1:URA3/+* |
| LY8673 | *MATa/α, Spc42-mCherry:kanMX6/+, $P_{TUB1}$GFP-TUB1:LEU2/+, cdc20::pCLB2-3HA-CDC20::KanMX6/cdc20::pCLB2-3HA-CDC20::KanMX6*, *cdc20pr-cdc20:URA3/ cdc20pr-cdc20:URA3* |
| LY8653 | *MATa/α, Spc42-mCherry:kanMX6/+, $P_{TUB1}$GFP-TUB1:LEU2/+, cdc20::pCLB2-3HA-CDC20::KanMX6/cdc20::pCLB2-3HA-CDC20::KanMX6*, *cdc20pr-cdc20S52A:URA3/ cdc20pr-cdc20S52A:URA3* |

| | |
|---|---|
| LY8654 | *MATa/α, Spc42-mCherry:kanMX6/+, $P_{TUB1}$GFP-TUB1:LEU2/+, cdc20::pCLB2-3HA-CDC20::KanMX6/cdc20::pCLB2-3HA-CDC20::KanMX6, cdc20pr-cdc20S127A:URA3/ cdc20pr-cdc20S127A:URA3* |
| LY8655 | *MATa/α, Spc42-mCherry:kanMX6/+, $P_{TUB1}$GFP-TUB1:LEU2/+, cdc20::pCLB2-3HA-CDC20::KanMX6/cdc20::pCLB2-3HA-CDC20::KanMX6, cdc20pr-cdc20S88AS89A:URA3/ cdc20pr-cdc20S88AS89A:URA3* |
| LY8656 | *MATa/α, Spc42-mCherry:kanMX6/+, $P_{TUB1}$GFP-TUB1:LEU2/+, cdc20::pCLB2-3HA-CDC20::KanMX6/cdc20::pCLB2-3HA-CDC20::KanMX6, cdc20pr-cdc20S127AT130AT131S132A:URA3/ cdc20pr-cdc20S127AT130AT131S132A:URA3* |
| LY5166 | *MATa/α, tor1-1::HIS3/tor1-1, fpr1::loxP-LEU2-loxP/fpr1::natMX4, RPL13A-2XFKBP12::TRP1/RPL13A-2XFKBP12::loxP, Spc42-mCherry:hphNT1/+, $P_{TUB1}$GFP-TUB1:URA3/+, ZIP1-GFP/+, Ipl1-FRB:kanMX6/ Ipl1-FRB:kanMX6* |
| LY8894 | *MATa/α, tor1-1::HIS3/tor1-1::HIS3, fpr1::natMX4/fpr1::natMX4, RPL13A-2XFKBP12::TRP1/RPL13A-2XFKBP12::TRP1, Spc42-mCherry:KanMX6/+, $P_{TUB1}$GFP-TUB1:LEU2/+, Ipl1-FRB:kanMX6/ Ipl1-FRB:kanMX6, cdc20::pCLB2-3HA-CDC20::KanMX6/cdc20::pCLB2-3HA-CDC20::KanMX6, cdc20pr-cdc20S88AS89A:URA3/ cdc20pr-cdc20S88AS89A:URA3* |
| LY3551 | *MATa/α, Spc42-mCherry:hphNT1/+, $P_{TUB1}$GFP-TUB1:URA3/$P_{TUB1}$GFP-TUB1:URA3, ZIP1-GFP/+, mad3::kanMX6/mad3::kanMX6* |
| LY4671 | *MATa/α, tor1-1::HIS3/tor1-1, fpr1::loxP-LEU2-loxP/fpr1::natMX4, RPL13A-2XFKBP12::TRP1/RPL13A-2XFKBP12::TRP1, Spc42-mCherry:hphNT1/+, $P_{TUB1}$GFP-TUB1:URA3/+, ZIP1-GFP/+, mad3::kanMX6/mad3::kanMX6* |

| | |
|---|---|
| LY7914 | *MATa/α, tor1-1/tor1-1, fpr1::loxP-LEU2-loxP/fpr1::natMX4, RPL13A-2XFKBP12::loxP/RPL13A-2XFKBP12::loxP, $P_{GAL1,10}$-NDT80:TRP1/ $P_{GAL1,10}$-NDT80:TRP1, Gal4-ER:URA3/Gal4-ER:URA3, Spc42-mCherry:hphNT1/+, $P_{TUB1}$GFP-TUB1:LEU2/ $P_{TUB1}$GFP-TUB1:LEU2, mad3::kanMX6/mad3::kanMX6, Ctf19-FRB:kanMX6/Ctf19-FRB:kanMX6* |
| LY9978 | *MATa/α, tor1-1::HIS3/tor1-1, fpr1::loxP-LEU2-loxP/fpr1::loxP-LEU2-loxP, RPL13A-2XFKBP12::TRP1/RPL13A-2XFKBP12::TRP1, $P_{GAL1,10}$-NDT80:TRP1/ $P_{GAL1,10}$-NDT80:TRP1, Gal4-ER:URA3/Gal4-ER:URA3, Spc42-mCherry:hphNT1/+, Ndc80-yeGFP:hphNT1/+, Ctf19-FRB:kanMX6/Ctf19-FRB:kanMX6* |
| LY9093 | *MATa/α, Spc42-mCherry:KanMX6/+, $P_{TUB1}$GFP-TUB1:LEU2/ $P_{TUB1}$GFP-TUB1:LEU2, cdc20::pCLB2-3HA-CDC20::KanMX6/cdc20::pCLB2-3HA-CDC20::KanMX6, cdc20pr-cdc20S88AS89A:URA3/ cdc20pr-cdc20S88AS89A:URA3, mad3::kanMX6/mad3::kanMX6* |
| LY9108 | *MATa, Spc42-mCherry:KanMX6, $P_{TUB1}$GFP-TUB1:LEU2, cdc20pr-cdc20:URA3, $P_{MET}$-HA3-CDC20:TRP1* |
| LY8691 | *MATa, tor1-1, fpr1::loxP-LEU2-loxP, RPL13A-2XFKBP12::loxP, ZIP1-GFP, mad3::kanMX6, Spc42-mCherry:hphNT1* |
| LY8961 | *MATa, Spc42-mCherry:KanMX6, $P_{TUB1}$GFP-TUB1:LEU2, cdc20pr-cdc20S127A:URA3, $P_{MET}$-HA3-CDC20:TRP1* |
| LY9061 | *MATa, Spc42-mCherry:KanMX6, $P_{TUB1}$GFP-TUB1:LEU2, cdc20pr-cdc20S52A:URA3, $P_{MET}$-HA3-CDC20:TRP1* |
| LY9095 | *MATa, Spc42-mCherry:KanMX6, $P_{TUB1}$GFP-TUB1:LEU2, cdc20pr-cdc20S88AS89A:URA3, $P_{MET}$-HA3-CDC20:TRP1* |

| | |
|---|---|
| LY9362 | *MATα*, *$P_{TUB1}$GFP-TUB1:LEU2*, *Spc42-mCherry:KanMX6*, *cdc20pr-cdc20:URA3*, *cdc20::NATMX4* |
| LY9480 | *MATα*, *$P_{TUB1}$GFP-TUB1:LEU2*, *Spc42-mCherry:KanMX6*, *cdc20pr-cdc20S127A:URA3*, *cdc20::NATMX4* |
| LY9449 | *MATα*, *$P_{TUB1}$GFP-TUB1:LEU2*, *Spc42-mCherry:KanMX6*, *cdc20pr-cdc20S88AS89A:URA3*, *cdc20::NATMX4* |
| LY3703 | *MATa/α, tor1-1::HIS3/tor1-1, fpr1::loxP-LEU2-loxP/fpr1:: natMX4, RPL13A-2XFKBP12::loxP/RPL13A-2XFKBP12::TRP1, Bub3-FRB:kanMX6/ Bub3-FRB:kanMX6, Spc42-mCherry:hphNT1/+, $P_{TUB1}$GFP-TUB1:URA3/$P_{TUB1}$GFP-TUB1:URA3, ZIP1-GFP/+* |
| LY9099 | *MATa/α, tor1-1::HIS3/tor1-1::HIS3, fpr1::natMX4/fpr1::natMX4, RPL13A-2XFKBP12::loxP/RPL13A-2XFKBP12::loxP, $P_{GAL1,10}$-NDT80:TRP1/ $P_{GAL1,10}$-NDT80:TRP1, Gal4-ER:URA3/Gal4-ER:URA3, Bub3-FRB:kanMX6/Bub3-FRB:kanMX6, Spc42-mCherry:hphNT1/+, $P_{TUB1}$GFP-TUB1:LEU2/+* |
| LY6932 | *MATa/α, Spc42-mCherry:KanMX6/+, $P_{TUB1}$GFP-TUB1:LEU2/$P_{TUB1}$GFP-TUB1:LEU2*, *alk1::natMX4/alk1::natMX4*, *alk2::hphNT1/alk2::hphNT1* |
| LY7031 | *MATa/α, tor1-1::HIS3/tor1-1::HIS3, fpr1::loxP-LEU2-loxP/fpr1::natMX4, RPL13A-2XFKBP12::loxP/RPL13A-2XFKBP12::loxP, Bub3-FRB:kanMX6/Bub3-FRB:kanMX6, Spc42-mCherry:kanMX6/+, $P_{TUB1}$GFP-TUB1:LEU2/$P_{TUB1}$GFP-TUB1:LEU2, alk1::natMX4/alk1::natMX4*, *alk2::hphNT1/alk2::hphNT1* |
| LY7730 | *MATa/α, Spc42-mCherry:KanMX6/+, $P_{TUB1}$GFP-TUB1:LEU2/+*, *alk1::natMX4/alk1::natMX4*, *alk2::hphNT1/alk2::hphNT1*, *mad3::kanMX6/mad3::kanMX6* |

| | |
|---|---|
| LY7575 | *MATa/α, Spc42-mCherry:KanMX6/+, $P_{TUB1}$GFP-TUB1:URA3/+, bir1::hphMX3:bir1-9xA-13xMyc:kanMX6:URA3/bir1::hphMX3:bir1-9xA-13xMyc:kanMX6:URA3* |
| LY7836 | *MATa/α, tor1-1::HIS3/tor1-1::HIS3, fpr1::natMX4/fpr1:: natMX4, RPL13A-2XFKBP12::loxP/RPL13A-2XFKBP12::loxP, Bub3-FRB:kanMX6/ Bub3-FRB:kanMX6, Spc42-mCherry:hphNT1/+, $P_{TUB1}$GFP-TUB1:URA3/$P_{TUB1}$GFP-TUB1:URA3, bir1::hphMX3:bir1-9xA-13xMyc:kanMX6:URA3/bir1::hphMX3:bir1-9xA-13xMyc:kanMX6:URA3* |
| LY7823 | *MATa/α, tor1-1::HIS3/tor1-1::HIS3, fpr1::natMX4/fpr1::natMX4, RPL13A-2XFKBP12::loxP/RPL13A-2XFKBP12::loxP, $P_{GAL1,10}$-NDT80:TRP1/ $P_{GAL1,10}$-NDT80:TRP1, Gal4-ER:URA3/Gal4-ER:URA3, Ctf19-FRB:kanMX6/Ctf19-FRB:kanMX6, Spc42-mCherry:hphNT1/+, $P_{TUB1}$GFP-TUB1:LEU2/$P_{TUB1}$GFP-TUB1:LEU2* |
| LY7805 | *MATa/α, tor1-1::HIS3/tor1-1::HIS3, fpr1::natMX4/fpr1::natMX4, RPL13A-2XFKBP12::loxP/RPL13A-2XFKBP12::loxP, $P_{GAL1,10}$-NDT80:TRP1/ $P_{GAL1,10}$-NDT80:TRP1, Gal4-ER:URA3/Gal4-ER:URA3, Ctf19-FRB:kanMX6/Ctf19-FRB:kanMX6, Bub3-FRB:kanMX6/Bub3-FRB:kanMX6, Spc42-mCherry:hphNT1/+, $P_{TUB1}$GFP-TUB1:LEU2/$P_{TUB1}$GFP-TUB1:LEU2* |
| LY7914 | *MATa/α, tor1-1/tor1-1, fpr1::loxP-LEU2-loxP /fpr1::natMX4, RPL13A-2XFKBP12::loxP/RPL13A-2XFKBP12::loxP, $P_{GAL1,10}$-NDT80:TRP1/ $P_{GAL1,10}$-NDT80:TRP1, Gal4-ER:URA3/Gal4-ER:URA3, Ctf19-FRB:kanMX6/Ctf19-FRB:kanMX6, mad3::kanMX6/mad3::kanMX6, Spc42-mCherry:hphNT1/+, $P_{TUB1}$GFP-TUB1:LEU2/$P_{TUB1}$GFP-TUB1:LEU2* |

| | |
|---|---|
| LY6695 | *MATa/α, tor1-1::HIS3/tor1-1, fpr1::natMX4/fpr1::natMX4, RPL13A-2XFKBP12::TRP1/RPL13A-2XFKBP12::TRP1, Spc105-FRB:kanMX6/Spc105-FRB:kanMX6,* $P_{REC8}$*-spc105*$^{RVAF}$*:LEU2/*$P_{REC8}$*-spc105*$^{RVAF}$*:LEU2, Spc42-mCherry:hphNT1/+,* $P_{TUB1}$*GFP-TUB1:URA3/*$P_{TUB1}$*GFP-TUB1:URA3* |
| LY7104 | *MATa/α, HTB2-mCherry:HIS3/+,* $P_{TUB1}$*GFP-TUB1:URA3/+, ZIP1-GFP/+* |
| LY7186 | *MATa/α, tor1-1::HIS3/tor1-1, fpr1::loxP-LEU2-loxP/fpr1::natMX4, RPL13A-2XFKBP12::loxP/RPL13A-2XFKBP12::loxP,* $P_{GAL1,10}$*-NDT80:TRP1/* $P_{GAL1,10}$*-NDT80:TRP1, Gal4-ER:URA3/Gal4-ER:URA3, HTB2-mCherry:HIS3/+* |
| LY3992 | *MATa/α, tor1-1::HIS3/tor1-1, fpr1::loxP-LEU2-loxP/fpr1::natMX4, RPL13A-2XFKBP12::loxP/RPL13A-2XFKBP12::TRP1, HTB2-mCherry:HIS3/+,* $P_{TUB1}$*GFP-TUB1:URA3/+, ZIP1-GFP/+* |
| LY5165 | *MATa/α, tor1-1::HIS3/tor1-1, fpr1::loxP-LEU2-loxP/fpr1::natMX4, RPL13A-2XFKBP12::loxP/RPL13A-2XFKBP12::TRP1, Ipl1-FRB:kanMX6/Ipl1-FRB:kanMX6, HTB2-mCherry:HIS3/+,* $P_{TUB1}$*GFP-TUB1:URA3/+, ZIP1-GFP/+* |
| LY3791 | *MATa/α, tor1-1::HIS3/tor1-1, fpr1::loxP-LEU2-loxP/fpr1::natMX4, RPL13A-2XFKBP12::loxP/RPL13A-2XFKBP12::TRP1, Bub3-FRB:kanMX6/Bub3-FRB:kanMX6, HTB2-mCherry:HIS3/+,* $P_{TUB1}$*GFP-TUB1:URA3/+, ZIP1-GFP/+* |
| LY6810 | *MATa/α, HTB2-mCherry:HIS3/+,* $P_{TUB1}$*GFP-TUB1:URA3/+, alk1::natMX4/alk1::natMX4, alk2::hphNT1/alk2::hphNT1* |
| LY6995 | *MATa/α, tor1-1::HIS3/tor1-1::HIS3, fpr1::natMX4/fpr1::natMX4, RPL13A-2XFKBP12::loxP/RPL13A-2XFKBP12::loxP, Bub3-FRB:kanMX6/Bub3-FRB:kanMX6, HTB2-mCherry:HIS3/+,* $P_{TUB1}$*GFP-TUB1:LEU2/+, alk1::natMX4/alk1::natMX4, alk2::hphNT1/alk2::hphNT1* |

| | |
|---|---|
| LY7556 | *MATa/α, tor1-1::HIS3/ tor1-1, fpr1::loxP-LEU2-loxP/fpr1::natMX4, RPL13A-2XFKBP12::TRP1/RPL13A-2XFKBP12::TRP1, $P_{GAL1,10}$-NDT80:TRP1/$P_{GAL1,10}$-NDT80:TRP1, Gal4-ER:URA3/Gal4-ER:URA3, Ctf19-FRB:kanMX6/Ctf19-FRB:kanMX6, HTB2-mCherry:HIS3/+* |
| LY7157 | *MATa/α, tor1-1::HIS3/ tor1-1::HIS3, fpr1::loxP-LEU2-loxP/fpr1::natMX4, RPL13A-2XFKBP12::loxP/RPL13A-2XFKBP12::loxP, $P_{GAL1,10}$-NDT80:TRP1/$P_{GAL1,10}$-NDT80:TRP1, Gal4-ER:URA3/Gal4-ER:URA3, Ipl1-FRB:kanMX6/Ipl1-FRB:kanMX6, HTB2-mCherry:HIS3/HTB2-mCherry:HIS3* |
| LY5391 | *MATa/α, tor1-1/tor1-1, fpr1::loxP-LEU2-loxP/fpr1::natMX4, RPL13A-2XFKBP12::loxP/RPL13A-2XFKBP12::loxP, $P_{GAL1,10}$-NDT80:TRP1/$P_{GAL1,10}$-NDT80:TRP1, Gal4-ER:URA3/Gal4-ER:URA3, Bub3-FRB:kanMX6/Bub3-FRB:kanMX6, HTB2-mCherry:HIS3/HTB2-mCherry:HIS3, ZIP1-GFP/+* |
| LY7868 | *MATa/α, tor1-1/tor1-1::HIS3, fpr1::loxP-LEU2-loxP/fpr1::natMX4, RPL13A-2XFKBP12::loxP/RPL13A-2XFKBP12::loxP, $P_{GAL1,10}$-NDT80:TRP1/$P_{GAL1,10}$-NDT80:TRP1, Gal4-ER:URA3/Gal4-ER:URA3, Bub3-FRB:kanMX6/Bub3-FRB:kanMX6, Ctf19-FRB:kanMX6/Ctf19-FRB:kanMX6, HTB2-mCherry:HIS3/+, $P_{TUB1}$GFP-TUB1:LEU2/+* |
| LY7610 | *MATa/α, HTB2-mCherry:HIS3/+, $P_{TUB1}$GFP-TUB1:URA3/ $P_{TUB1}$GFP-TUB1:URA3, bir1::hphMX3:bir1-9xA-13xMyc:kanMX6:URA3/bir1::hphMX3:bir1-9xA-13xMyc:kanMX6:URA3* |
| LY7955 | *MATa/α, tor1-1::HIS3/tor1-1::HIS3, fpr1::natMX4/fpr1::natMX4, RPL13A-2XFKBP12::loxP/RPL13A-2XFKBP12::loxP, Bub3-FRB:kanMX6/Bub3-FRB:kanMX6, bir1::hphMX3:bir1-9xA-13xMyc:kanMX6:URA3/bir1::hphMX3:bir1-9xA-13xMyc:kanMX6:URA3, HTB2-* |

| | |
|---|---|
| | *mCherry:HIS3/HTB2-mCherry:HIS3,* $P_{TUB1}$*GFP-TUB1:URA3/* $P_{TUB1}$*GFP-TUB1:URA3* |

**Table S2. Reagents used in this study**

| Reagent | Source | Identifier |
|---|---|---|
| Yeast extract | Thermo Fisher Scientific | Ref#212720, Lot#2352585 |
| Peptone | Thermo Fisher Scientific | Ref#211820, Lot#1263496 |
| Dextrose (D-Glucose) Anhydrous | Fisher Chemical | Lot#204841 |
| Potassium acetate | Fisher Bioreagents | Lot#210889 |
| Yeast nitrogen base without amino acids | Thermo Fisher Scientific | Ref#291920, Lot#9148845 |
| Synthetic complete mixture drop-out: Complete | Formedium | Ref# DSCK2500, Batch# FM0A416/006650 |
| Bactoagar | Thermo Fisher Scientific | Cas#: 214010 |
| β-estradiol | Sigma | Cas#: 50-28-2 |
| Rapamycin | Fisher BioReagents | Cas#: 53123-88-9 |
| Alpha factor | Zymo research | Cas#: Y1001 |
| Concanavalin | Sigma | Cas#: 11028-71-0 |
| Acetone | VWR | Cas#:67-64-1; Lot#: 20L0956101 |

| | | |
|---|---|---|
| Paraformaldehyde (PFA) | Macron Fine Chemicals | Cas#: 30525-89-4 |
| Glycine | Sigma | Cat#G7126-1KG, LOT# SLCH8988 |
| MEM with Earle's Salts | Sigma | Cat# M0268-10x1L |
| Pyruvate (Sodium Salt) | Sigma | Cat# P4562-25G |
| Gentamycin | Sigma | Cat# G1272-10ML |
| Hepes | Sigma | Cat# h3784-100G |
| PVP | Sigma | Cat# P2307-100G |
| BSA | Sigma | Cat# A4503 |
| Tween-20 | Sigma | Cat# 274348 |
| Sodium azide ($NaN_3$) | Sigma | Cat# 71289 |
| NaCl | Sigma | Cat# S5886-500G |
| KCl | Sigma | Cat# P5405-250G |
| $KH_2PO_4$ | Sigma | Cat# P5655-100G |
| $MgSO_4$ $7H_2O$ | Sigma | Cat# M7774-500G |
| Pyruvic acid (sodium salt) | Sigma | Cat# P4562-25G |
| $CaCl_2$ $2H_2O$ | Sigma | Cat# C7902-500G |
| DL-Lactic acid (sodium salt, 60% syrup) | Sigma | Cat#: L900-100ML |
| Taurine | Sigma | Cat# T0625-10G |
| EDTA | Sigma | Cat# E5134-100G |
| $NaHCO_3$ | Sigma | Cat# S5761-500G |
| Gentamicin | Sigma | Cat# G1272-10ML |
| Phenol red | Sigma | Cat# P5530-5G |

| | | |
|---|---|---|
| BSA | Sigma | Cat# A4503-100G |
| Milrinone | Sigma | Cat#: M4659 |
| L-Glutamine | Sigma | SLBS6549 |
| Vectashield | Vector Laboratories | Cat#: H-1000 |
| 6-diamidino-2-phenylindole, Dihydrochloride (DAPI) | Thermo Fisher Scientific | Cat#D1306, 2301042 |
| Dimethyl sulfoxide (DMSO) | Sigma | Cat#472301 |
| Ethanol | Fisher | Cas#64-17-5 |
| 0.5mm glass beads | BioSpec Products | Cat# 11079105 |
| Tyrode | Millipore Sigma | MR-004-D |

**Table S3. Drugs used for mouse work**

| Drug | Source | Identifies |
|---|---|---|
| 5-Iodtubercidin (5-Itu) | Cayman Chemical | Cas#24386-93-4, Item#: 10010375 |
| BAY-1816032 | MedChem Express | Cat#: HY-103020/CS-0023128 (10mM*1mL in DMSO) |
| ZM447439 | Tocris Bioscience | Cas#: 331771-20-1, Cat#: 2458 |

**Table S4. Antibodies used for mouse work**

| Antibody | Source | Identifier |
|---|---|---|
| Rabbit monoclonal anti-α-tubulin Alexa Fluor 488 conjugated | Cell Signaling | Cas#2125s; RRID:AB_2619646 |

| | | |
|---|---|---|
| Human polyclonal anti-CREST/ACA | Antibodies Incorporated | Cat#: 15-234; RRID:AB_2687472 |
| Goat anti-human Alexa-Fluor 633 | Life Technologies | Cat#A21091; RRID:AB_2535747 |
| Rabbit anti-mAurC Antibody | FORTIS | A300-BL1217 |
| Alexa Fluor 568 donkey anti-rabbit IgG (H+L) | Invitrogen | A10042 |
| Histone H2AT120ph pAb | Active Motif | Cat#39391 |
| Goat anti-Rabbit IgG Alexa-Fluor 488 | Life Technologies | Cat#A11034 |

**Table S5. Programs used for image acquisition and analysis**

| Program | Source |
|---|---|
| NIS-Elements software | Nikon |
| ImageJ | NIH, https://imagej.nih.gov |
| Graphpad Prism | https://www.graphpad.com/scientific-software/prism/ |
| Adobe Illustrator | https://www.adobe.com/products/illustrator.html |
